# Stiffness-dependent active wetting enables optimal collective cell durotaxis

**DOI:** 10.1101/2022.07.24.501310

**Authors:** Macià-Esteve Pallarès, Irina Pi-Jaumà, Isabela Corina Fortunato, Valeria Grazu, Manuel Gómez-González, Pere Roca-Cusachs, Jesus M de la Fuente, Ricard Alert, Raimon Sunyer, Jaume Casademunt, Xavier Trepat

## Abstract

The directed migration of cellular clusters enables morphogenesis, wound healing, and collective cancer invasion. Gradients of substrate stiffness are known to direct the migration of cellular clusters in a process called collective durotaxis, but underlying mechanisms remain unclear. Here, we unveil a connection between collective durotaxis and the wetting properties of cellular clusters. We show that clusters of cancer cells dewet soft substrates and wet stiff ones. At intermediate stiffness, at the crossover from low to high wettability, clusters on uniform-stiffness substrates become maximally motile, and clusters on stiffness gradients exhibit optimal durotaxis. Durotactic velocity increases with cluster size, stiffness gradient, and actomyosin activity. We demonstrate this behavior on substrates coated with the cell-cell adhesion protein E-cadherin and then establish its generality on substrates coated with extracellular matrix. We develop a physical model of three-dimensional active wetting that explains this mode of collective durotaxis in terms of a balance between in-plane active traction and tissue contractility, and out-of-plane surface tension. Finally, we show that the distribution of cluster displacements has a heavy tail, with infrequent but large cellular hops that contribute to durotactic migration. Our study demonstrates a physical mechanism of collective durotaxis, through both cell-cell and cell-substrate adhesion ligands, based on the wetting properties of active droplets.

## Introduction

The ability of cells to migrate following gradients of environmental properties drives numerous biological processes in health and disease^1^. Cells undergo directed migration in response to gradients of chemical factors^2^, electric fields^3^, and mechanical properties of their microenvironment^4–7^. This latter process, called durotaxis, has been implicated in morphogenesis^8,9^, wound healing^10^ and cancer invasion^11^.

Durotaxis was originally discovered as the directed migration of single cells following a gradient in the stiffness of an underlying substrate^4^. Later on, cell collectives were found to perform durotaxis much more efficiently than single cells^12,13^. Collective durotaxis has now been established *in vitro* as the asymmetric spreading of an epithelial monolayer^12,13^ and *in vivo* as the key mechanism driving the cohesive migration of the neural crest cluster during development of *Xenopus laevis*^9^. Despite the growing recognition of the physiological relevance of durotaxis in development and disease, underlying mechanisms remain poorly understood. This limitation arises, in part, from the lack of data on how durotaxis depends on key physical variables of the motile cluster, such as its size, adhesion, threedimensional shape, and active forces. Moreover, current theoretical models^12,14–21^ do not integrate all these physical variables into a physical framework that explains collective durotaxis.

Durotaxis of single cells and clusters has so far been studied in the presence of gradients in the stiffness of the extracellular matrix (ECM)^4–7,12,22,23^. However, important migratory processes during development and cancer progression take place in contexts lacking ECM^24,25^. This is illustrated by border cell migration during *Drosophila* oogenesis, which is mediated by E-cadherin-based adhesion between the migratory cluster and surrounding stationary cells^25,26^. Other examples of cell migration through cadherin adhesion include zebrafish primordial germ cells^27^, mouse retinal endothelial cells^28^, and mouse neuronal precursors^29^. Cadherin-based migration is also likely to play a role in epithelial tumors in which collective invasion and remodeling are predominantly governed by cell-cell adhesion rather than by the ECM^30^. Although collective migration through cadherin receptors is well established, whether these cell-cell adhesion proteins can mediate durotaxis is unknown.

Here we provide a systematic mechanical analysis of collective durotaxis in three-dimensional epithelial clusters. We begin studying durotaxis on substrates coated with E-cadherin. We show that clusters of human epidermal carcinoma cells (A431) display low motility on the soft and stiff regions of the substrate, where they fully dewet and wet the surface, respectively. At intermediate stiffness, close to the crossover between low and high wettability, cell clusters are maximally motile on uniform-stiffness substrates and exhibit optimal durotaxis on stiffness gradients. We develop a continuum model of threedimensional active wetting that explains this non-monotonic durotactic behavior in terms of a balance between cellular traction, contractility and surface tension. Cell clusters then perform cohesive durotactic migration as their interface advances on the stiff side and retracts from the soft side. We show that this physical mechanism applies not only to surfaces coated with cell-cell adhesion ligands but also with cell-ECM ligands as long as clusters are brought close to the crossover between low and high wettability, which we call the neutral wetting regime.

## Results

### Cell clusters become most motile at their neutral wetting regime

We first studied the spontaneous migration of cell clusters (83±54 cells/cluster, mean + SD, Extended Data Fig. 1a) on polyacrylamide gels of uniform stiffness (0.2, 6, 24 and 200 kPa) functionalized with oriented E-cadherin extracellular domains (Fig. 1; Extended Data Fig. 2). Using a custom-made tracking algorithm, we measured the cluster position for 14h at 10 min intervals (Fig. 1a-c; Supplementary video 1). The dependence of cluster velocity with stiffness was non-monotonic; it was minimal at low stiffness (0.2 kPa), peaked at intermediate stiffness (24 kPa), and then decreased at high stiffness (200 kPa) (Fig. 1b,d). This non-monotonic behavior coincided with different regimes of cluster spreading, which we interpret within the conceptual framework of tissue wetting^14,15,31–38^. At low stiffness, clusters were nearly spherical and the contact angle (*θ*) between the cluster and the substrate was close to 180°, indicating full dewetting (i.e. complete retraction) (Fig. 1a,e,f). By contrast, at high stiffness, clusters spread to form a monolayered epithelium with a low contact angle, indicating full wetting (i.e. complete spreading). At intermediate stiffness, for contact angles around 90°, clusters displayed highly dynamic protrusions (Fig. 1f; Supplementary video 1,2). To characterize these protrusions, we imaged clusters transfected with mCherry-Lifeact at high spatial and temporal resolution. The resulting movies revealed that these protrusions are dynamic filopodia (Supplementary video 3), in which actin flows retrogradely towards the cluster core (Supplementary video 4). Taken together, these data suggest that at the crossover between low and high wettability (*θ*~90°), which we define as the neutral wetting regime, clusters become maximally motile by rapidly engaging and disengaging actin-rich protrusions with the substrate.

**Figure 1.**
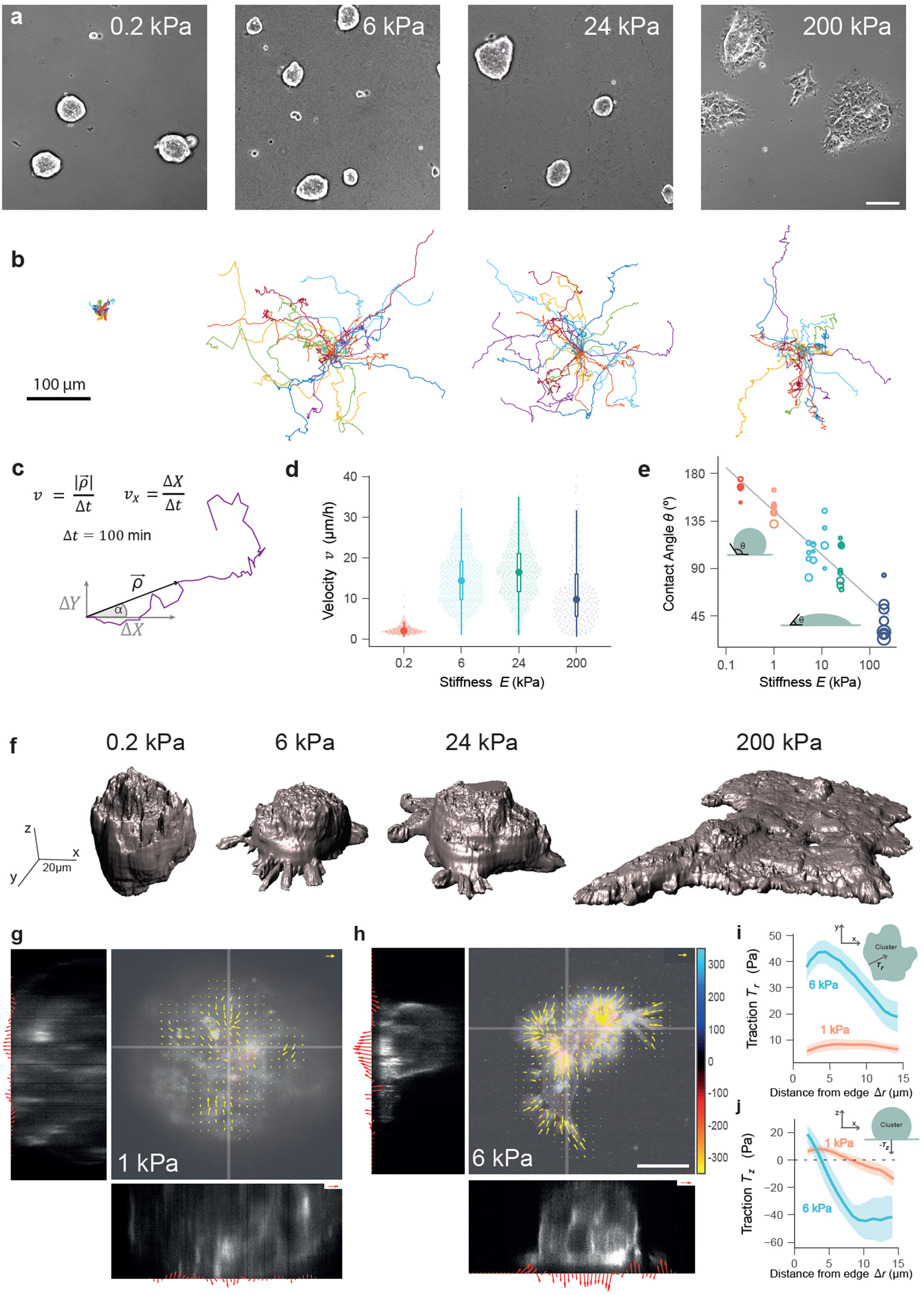
Epithelial cell clusters on E-cadherin-coated substrates show optimal motility in the neutral wetting regime. **a**, Representative phase contrast images of A431 cell clusters seeded on E-cadherin-coated gels of uniform stiffness of 0.2, 6, 24 and 200 kPa. Scale bar, 100 μm. **b**, Representative trajectories of clusters migrating on the E-cadherin-coated gels shown in panel (a). **c**, Scheme representing the workflow to track cluster trajectories and velocity. Velocity was computed at 100 min intervals. **d**, Cluster velocity at different substrate stiffness. Small dots represent individual clusters. The central dot in front of the boxplot represents the median. For the sake of visualization, clusters above the percentile 99.5% are not represented. Permutation tests (two tailed) were performed, in which significant differences were observed across all combinations (p-values < 0.0001). n= 296 clusters (0.2 kPa), n= 646 clusters (6 kPa), n= 561clusters (24 kPa), n= 266 clusters (200 kPa). **e**, Contact angle *θ* between the cell cluster and the substrate for different substrate stiffness. Each dot is the averaged contact angle for one cluster. Circle size is proportional to cluster average diameter. n= 43 clusters. **f**, 3D rendering of representative clusters on substrates of different stiffness. **g-h**, Traction forces exerted by representative clusters on 1 and 6 kPa E-cadherin-coated gels. Yellow vectors represent traction forces in the *xy*-plane while red vectors represent traction forces projected on the corresponding lateral planes (*xz* and *yz*) along the gray lines shown in the central panels (reference vectors are 50 Pa, scale bar is 25 μm). **i**, Mean of the radial component of the traction force in the *xy*-plane for 1 kPa (n= 35 clusters) and 6 kPa gels (n= 19 clusters) as a function of distance from the cluster edge. **j**, Mean of the vertical component of the traction force for 1 kPa (n= 13 clusters) and 6 kPa gels (n= 7 clusters) as a function of distance from the cluster edge. Shaded areas in i-j represent 95% confidence interval (CI).

We next examined whether these local protrusions generated traction forces. We performed 3D Traction Force Microscopy (TFM) experiments on clusters seeded on polyacrylamide gels of 1 and 6 kPa in stiffness (Fig. 1g-j). At higher stiffness, TFM showed insufficient resolution to robustly measure the three traction components. We characterized traction profiles through their radial component (*T_r_*), defined as the component perpendicular to the cluster edge in the substrate plane, and through the normal component to the substrate (*T_z_*). Both on 1 kPa and 6 kPa substrates, radial tractions pointed towards the center of the cluster (Fig. 1g-i). Normal tractions were positive near the cluster edges and became negative towards the cluster center. These data reveal a cluster surface tension (*γ*) that pulls the cluster edge upwards at the contact line with the substrate. Consistent with this picture, cell protrusions at the cluster edge were not parallel to the substrate but rather formed acute angles with it (Fig. 1f; Extended Data Fig. 3). Upwards traction at the cluster edge is balanced by a pressure that pushes the cluster core into the substrate. Whereas the spatial profiles of *T_z_* and *T_r_* displayed qualitative similarities on 1 kPa and 6 kPa, the magnitude of both components increased with stiffness, indicating that in-plane tractions and surface tension are mechanosensitive^39–42^ (Fig. 1i,j).

### Cell clusters exhibit durotaxis on E-cadherin substrates

The highly dynamic state of epithelial clusters in the neutral wetting regime led us to hypothesize that they might be particularly responsive to gradients in substrate stiffness. To test this hypothesis, we fabricated substrates exhibiting stiffness gradients^43^ and functionalized them with the oriented extracellular domain of E-cadherin at uniform density (see Methods and Extended Data Fig. 4). As in previous experiments, we tracked the migration of cell clusters for 14h at 10 min intervals (Fig. 2a; Supplementary videos 5,6). After each experiment, we measured the substrate stiffness profile with Atomic Force Microscopy (AFM). Using this approach, we built a large dataset matching the local mechanical properties of the substrate with the instantaneous velocity of each cluster. Clusters migrating on stiffness gradients showed a significantly positive velocity (*v_X_*) along the direction of the gradient, indicating durotaxis towards increasing stiffness (Fig. 2b). This biased migration was also evident when comparing the angular distribution of cluster trajectories for uniform (Fig. 2c) and gradient (Fig. 2d) gels. These experiments show that durotaxis is not restricted to integrin-mediated migration on ECM substrates. Rather, the cell migration machinery can also drive durotactic responses through cadherin receptors.

**Figure 2.**
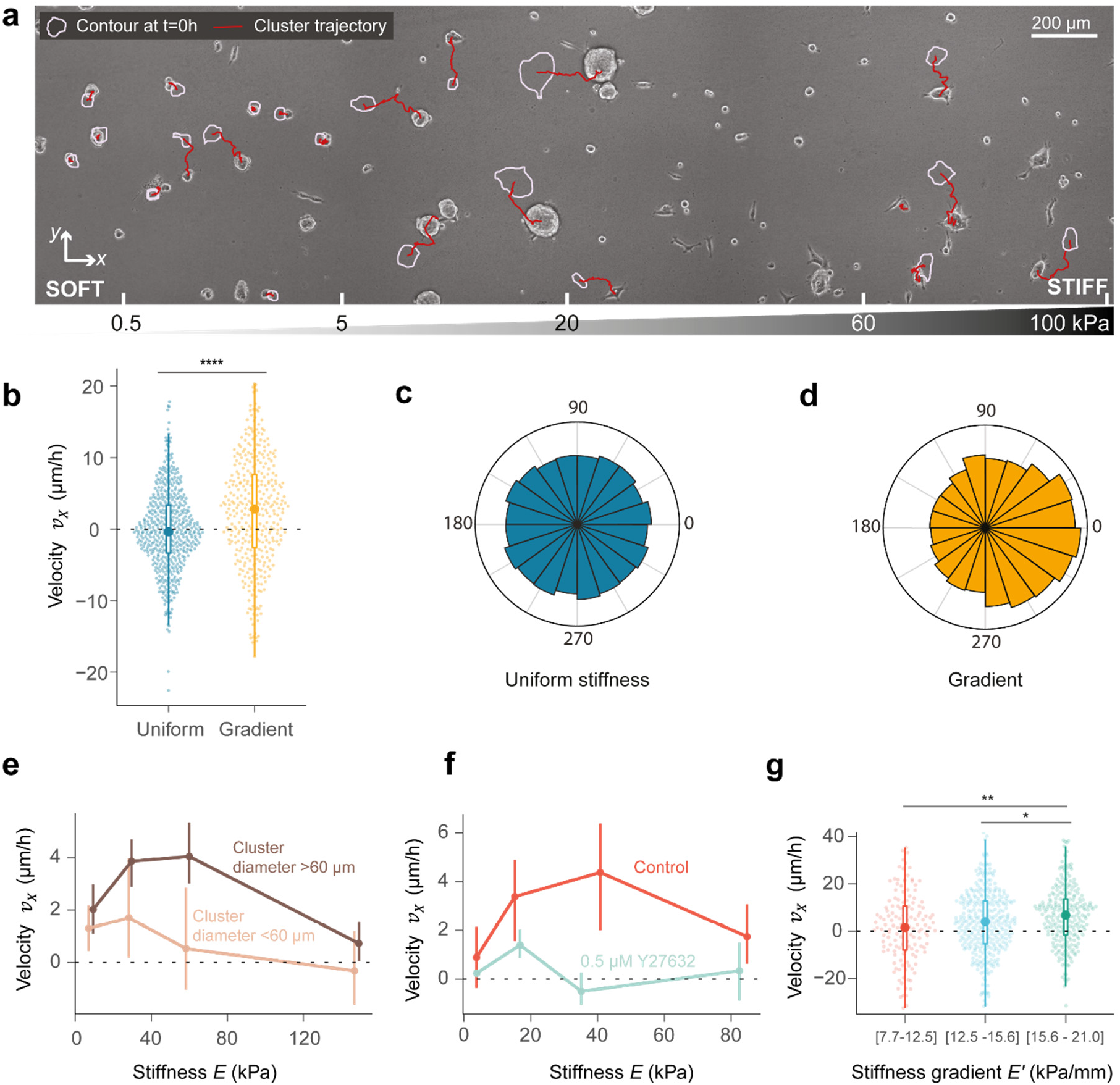
Collective durotaxis is optimal in the neutral wetting regime. **a**, Representative phase-contrast image of A431 cell clusters migrating on a stiffness gradient coated with E-cadherin. Image was taken at 10h. The original position (0h) of each cluster is represented by a purple outline. The red line represents the trajectory obtained by time-lapse microscopy. The bottom scale indicates the stiffness at each point of the image. **b**, Cluster velocity *v_X_* along the direction of the gradient on uniform (6 kPa) and gradient stiffness gels. Small dots represent individual clusters. The central dot in front of the boxplot represents the median. (Permutation t-test (two tailed), p < 0.0001). n= 527 clusters (Uniform), n= 366 clusters (Gradient, data pooled from all regions of the substrate). **c-d**, Distribution of the angle *a* between the instantaneous velocity vector and the *x*-axis (see Fig. 1c) for uniform stiffness gels (c) and stiffness gradients (d). **e**, Durotactic velocity as a function of stiffness. Light brown labels small clusters (diameter < 60 μm), whereas dark brown labels large clusters (diameter > 60 μm). Vertical bars indicate 95% CI. Bin 1 (0 < *E* ≤ 20 kPa): diameter < 60 μm, n=72 clusters (529 displacements); diameter > 60 μm, n=90 clusters (584 displacements). Bin 2 (20 < *E* ≤ 40 kPa): diameter < 60 μm, n=30 clusters (136 displacements); diameter > 60 μm, n=86 clusters (549 displacements). Bin 3 (40 < *E* ≤ 80 kPa): diameter < 60 μm, n=37 clusters (205 displacements); diameter > 60 μm, n=106 clusters (629 displacements). Bin 4 (*E* > 80 kPa): diameter < 60 μm, n=53 clusters (248 displacements); diameter > 60 μm, n=186 clusters (1103 displacements). **f**, Durotactic velocity in control clusters (red) and in clusters treated with Y27632 (0.5 μM, light blue). Data includes only clusters > 60 μm. Vertical bars represent 95% CI. Bin 1 (0 < *E* ≤ 10 kPa): control, n=11 clusters (82 displacements); 0.5 μM Y27632, n=28 clusters (216 displacements). Bin 2 (10 < *E* ≤ 25 kPa): control: n=19 clusters (105 displacements); 0.5 μM Y27632, n=29 clusters (164 displacements). Bin 3 (25 < *E* ≤ 60 kPa): control, n=14 clusters (77 displacements); 0.5 μM Y27632, n=11 clusters (69 displacements). Bin 4 (*E* > 60 kPa): control, n=13 clusters (71 displacements); 0.5 μM Y27632, n=18 clusters (70 displacements). **g**, Durotactic velocity for different stiffness gradients for a fixed starting stiffness *E*=18 ± 5 kPa. Clusters on steeper gradients showed significantly more durotaxis than those on milder ones (Permutation test, p-values: Bin 1 vs Bin 2, 0.1014; Bin 2 vs Bin 3, 0.0463; Bin 1 vs Bin 3, 0.0026). Each point represents a displacement. The central dot in front of the boxplot represents the median. For the sake of visualization, clusters above the percentile 99.5% are not represented. Bin 1 (7.7 < *E’* ≤ 12.5 kPa/mm): n=20 clusters (141 displacements); Bin 2 (12.5 < *E’* ≤ 15.6 kPa/mm): n=45 clusters (339 displacements); Bin 3 (15.6 < *E’* ≤ 21 kPa/mm): n=41 clusters (248 displacements).

### Durotaxis depends on stiffness, stiffness gradient, cluster size and cell contractility

We next explored whether durotaxis depends on local substrate stiffness (E). As in the case of substrates of uniform stiffness, clusters dewetted regions of low stiffness and wetted those of high stiffness (Fig. 2a; Supplementary video 5). In these two extreme cases, the cluster durotactic velocity *v_X_* was low (Fig. 2e). However, in regions of intermediate stiffness, clusters were in the neutral wetting regime and durotaxis peaked. Thus, as in uniform substrates (Fig. 1d), cluster velocity was maximal at intermediate stiffness, but in this case migration was directed towards higher stiffness rather than random.

To characterize durotaxis, we studied the role of cluster size, cell contractility, and stiffness gradient. We found that large clusters (diameter >60 μm) were more durotactic than smaller ones (Fig. 2e), and their velocity peaked at higher stiffness (Fig. 2e). To decrease cell contractility, we treated cells with a low dose (0.5 μM) of the ROCK inhibitor Y-27632. As expected, for any given stiffness this treatment resulted systematically in smaller contact angles (Extended Data Fig. 1b). We also found that the decrease in contractility reduces durotaxis and shifts the durotaxis peak to lower stiffness compared to untreated clusters (Fig. 2f; Supplementary video 7). Finally, to study how durotaxis depends on the stiffness gradient, we generated substrates with low, middle, and high steepness. Since durotaxis varies with local stiffness, we measured cluster velocity on each gradient substrate at a similar starting stiffness (18 ± 5 kPa). We found a significant increase in durotaxis with the stiffness gradient (Fig. 2g). Together, these results establish that durotaxis is optimal in the neutral wetting regime and depends on cluster size, contractility, and stiffness gradient.

### A three-dimensional model of active wetting explains non-monotonic tissue durotaxis

So far, our data show that the peaks in cluster velocity and durotaxis correlate with the wetting state of the cell clusters. To understand how the tissue wetting properties might lead to collective durotaxis, we model clusters as active fluid droplets that partially wet the substrate. Accordingly, we describe a cluster as a spherical cap of radius *R*_sphere_, whose contact surface with the substrate is a circular cell monolayer of radius *R* (Fig. 3a). Based on the observation that protrusive activity is largely restricted to cells in contact with the substrate (Extended Data Fig. 3, Supplementary Videos 2-4), we assume that the dynamics of the droplet is controlled by the in-plane forces in the basal monolayer, which we model as a 2D active polar fluid, extending previous work^14,15,31,38,44^. The cells at the periphery of this monolayer are polarized outwards and exert two types of active forces (Fig. 3a, inset): cell-substrate traction with a maximum value *ζ_i_*, which promotes tissue spreading, and cell-cell contractility with magnitude *ζ* < 0, which promotes tissue retraction. This cell-cell contractility refers to active contractile stresses within and between cells, generated by the actomyosin cytoskeleton and transmitted across the cell monolayer through cell-cell junctions. Previous work showed that the competition between traction and contractility in this 2D model gives rise to an active wetting transition between monolayer spreading (wetting) and retraction (dewetting)^14,31,38^.

**Figure 3.**
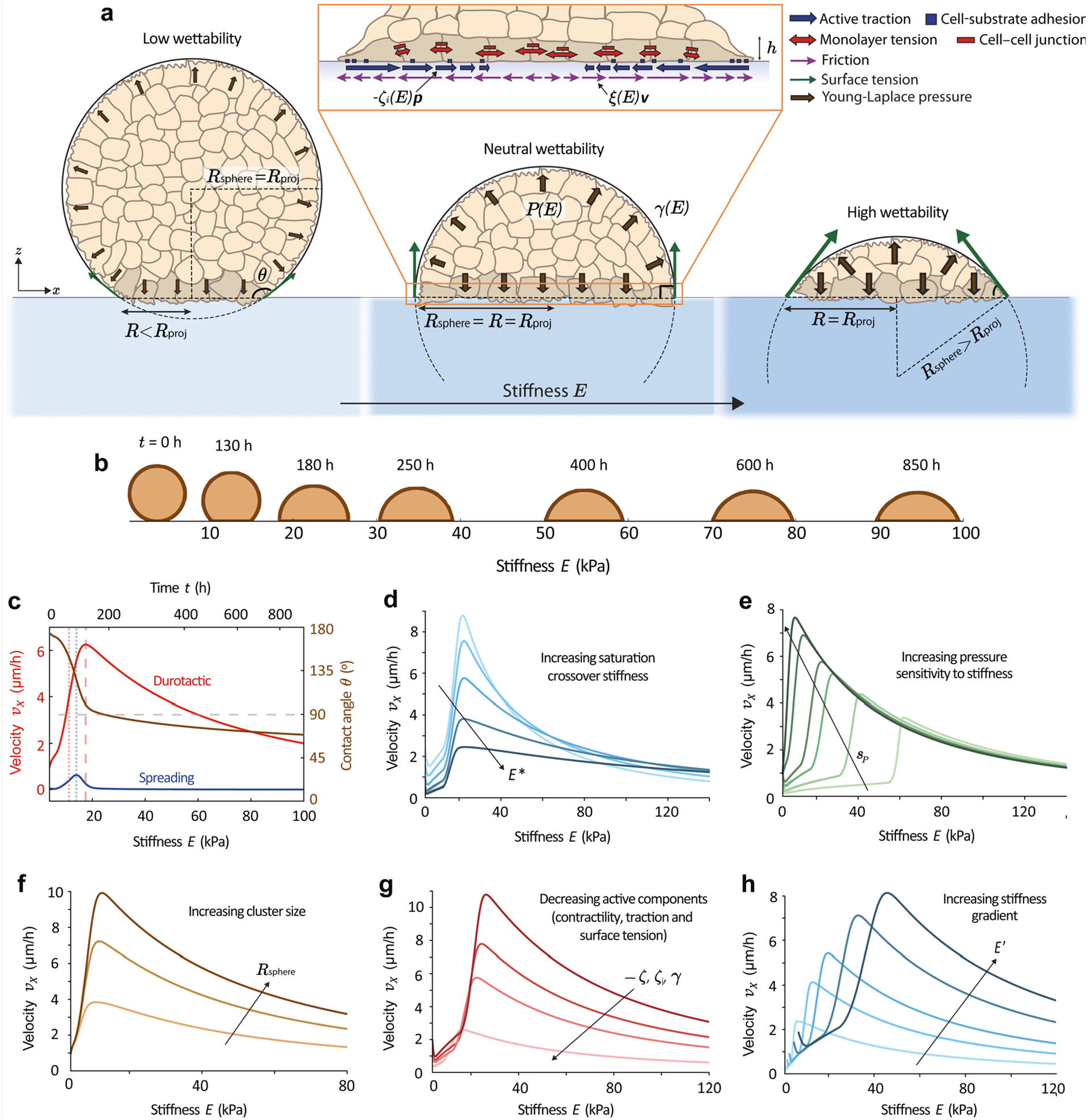
A three-dimensional model of active wetting explains non-monotonic tissue durotaxis. **a**, Scheme of the model for a cell cluster on substrates of different stiffness *E*. *R*_proj_ is the projected radius of the cell cluster, which is the radius measured in phase-contrast images. The inset shows a zoom-in of the basal cell monolayer (represented with darker cells for visualization) with schematic representations of the cell-substrate forces and the monolayer tension. In all panels **b**-**h**, the initial contact radius is *R*_0_ = 3.0 μm and height *H*_0_ = 56.8 μm (giving a contact angle of *θ*_0_ = 174° and volume *V* = 92500*π*/3 μm^3^), and the initial substrate stiffness is *E*_0_ = 3.8 kPa. Except for the parameters that we are changing in each panel, other parameter values are listed in Table SI, with a simulation time-step of Δ*t* = 6 min. See simulation details in the Methods section. **b-c,** Representative example of the velocity and shape dynamics of a migrating cluster with a constant volume, showing the snapshots of the cluster shape (**b**) and the non-monotonic dependence of the durotactic velocity *v_X_* with stiffness (in red), the spreading velocity (in blue) and the decrease of the contact angle (in brown) (**c**). Since the cluster migrates towards stiffer regions of the substrate, the stiffness and time axes run in parallel. In this case, we have taken a saturating pressure profile with stiffness to study how the angle keeps decreasing at large stiffness, with *P*^∞^ = 0.6 kPa and 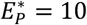 and following the same expressions than in Eq. (S12). **d,** The saturation crossover stiffness *E*\* controls the decrease of velocity at high stiffness as it tunes both friction and active traction increase and saturation with stiffness. Here, *E*\* = 50,80,140,260,450 kPa from lighter to darker blue. **e**, The increase of surface tension, and hence pressure, with stiffness controls the increase of velocity at low stiffness. Here, *P*′ = (0.1,0.2,0.4,0.6,1.5,3.0) Pa/μm, which corresponds to *s_P_* = (0.3,0.6,1.2,1.8,4.5,9.1) ·10^-2^, from lighter to darker green. **f,** Larger clusters are more durotactic and feature the velocity peak at higher stiffness. Here, the initial sizes are *R*_sphere_ = 13.2,28.5,61.4 μm (from lighter to darker brown), giving volumes of 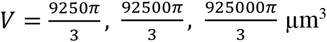, and *γ*(*E*) = 0.05 + 0.03(*E* – *E*_0_)/*E*′ mN/m for all sizes. **g,** Clusters with reduced active forces present a lower durotactic velocity with its peak shifted to lower stiffness. Here, contractility and traction are multiplied by factors *α_ζ_* = *α_ζ_i__* = 0.4,1,1.4,2, and surface tension by *α_γ_ =* 0.7,1,1.2,1.5 (smaller factors towards lighter red). **h,** Larger stiffness gradients produce an increase of durotactic velocity and a displacement of its peak towards stiffer regions. Here, *E*′ = 10,20,30,50,70 kPa/mm (from lighter to darker blue).

Here, we extend the theory of active wetting to 3D droplets (Supplementary Note). We propose a generalized Young-Dupré force balance between the active forces in the basal monolayer and the out-of-plane surface tension *γ* of the cell cluster, defining a contact angle *θ* (Fig. 3a). The horizontal component of the surface tension, −*γ* cos *θ*, combines with the monolayer active forces to drive either spreading or retraction, damped by both monolayer viscosity *η* and substrate friction *ξ*. For sufficiently large surface tension, the tissue may reach a stable equilibrium with a static contact angle defining partial wetting with high (*θ* < 90°) or low (*θ* > 90°) wettability (Supplementary Note). In turn, the vertical component of surface tension is balanced by the Young-Laplace pressure *P* = 2*γ*/*R*_sphere_ exerted on the contact surface (Fig. 3a). Assuming that the surface tension, and hence the Laplace pressure, is uniform across a cluster, we use this relation to infer the value of *γ* in our experiments from measured vertical traction forces (Fig. 1g-j), which provide a direct measurement of *P* (see Table 2 in Methods).

To capture collective durotaxis, we take into account that cellular forces depend on substrate stiffness. Following previous work^14,45–50^, we assume that both the active traction *ζ_i_* and the friction coefficient *ξ* increase and saturate with the substrate’s Young modulus *E*. This is consistent with our measurements, which show that radial in-plane tractions increase with substrate stiffness (Fig. 1g-i). In addition, our measurements reveal that out-of-plane tractions also increase with stiffness (Fig. 1j), implying that the tissue surface tension features an active mechanosensitive response^51,52^. Consistently, we assume that the pressure *P* is an increasing function of *E*, which we take as linear for simplicity. Altogether,

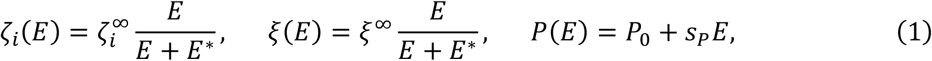

where 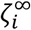 and *ξ*^∞^ are saturation values, *E*\* is a characteristic stiffness of force saturation, *P*_0_ is the bare pressure, and *s_P_* is the pressure sensitivity to stiffness.

With this model, we simulate the motion of a cluster up a stiffness gradient *E*(*x*) = *E*_0_ + *E*′*x* (Fig. 3b). As the cluster moves to stiffer regions, its contact angle decreases (Fig. 3c, brown), but its durotactic velocity varies non-monotonically (Fig. 3c, red), in agreement with our experimental measurements (Fig. 2e,f). We explain the origin of this non-monotonic behavior below.

Irrespective of the wettability of the 3D droplet, the balance of in-plane forces at the basal monolayer yields two basic predictions. First, as predicted in Refs.^14,15^, the durotactic velocity *v_X_* increases with the difference between the active traction at the front (stiff edge) and at the rear (soft edge) of the cluster (Supplementary Note). Hence, *v_X_* increases with the stiffness gradient, consistently with our measurements (Fig. 2g). Second, at high stiffness, active traction forces saturate. Hence, the active traction difference across the cluster decreases, which results in slower durotaxis (Fig. 3d). In parallel, the friction coefficient increases (and also saturates) with stiffness, which also results in slower durotaxis (Supplementary Note). This slowdown was also observed in our experimental measurements (Fig. 2e,f), and it can arise either at *E* > *E*\* due to traction saturation or at *E* < *E*\* due to increasing friction (Fig. 3d).

However, our experiments also show that the durotactic velocity increases with stiffness in the low stiffness area of the substrate (Fig. 2e,f). This increase could be explained by the fact that the stiffness gradient generated by our fabrication method increases in this region of the substrate (Extended Data Fig. 4a). To test this possibility, we took advantage of the gel-to-gel variability (Extended Data Fig. 4b) and, for each stiffness bin, we restricted our analysis to the data points with a gradient of 40±10 kPa/mm. We found that in these conditions of nearly constant gradient, the non-monotonic behavior of durotactic velocity with stiffness was retained (Extended Data Fig. 5). This result indicates that the sharp increase in durotactic velocity in the low stiffness region of the gel cannot be attributed to changes in the stiffness gradient.

Instead, using our 3D active wetting theory, we predict that this feature arises from the wettability of the tissue, i.e. the contact angle *θ* determined by the generalized Young-Dupré force balance (Supplementary Note). As in our experiments (Fig. 1e,f), at low stiffness the tissue has high contact angles (*θ* > 90°, Fig. 3b,c). As a result, the contact radius *R* is small. This leads to a small active traction difference across the tissue, and hence a small durotactic velocity. However, the contact angle is large, and therefore surface tension pulls the cluster edges out (Fig. 3a, left) and gives rise to a positive spreading velocity (Fig. 3c, blue). As a result, the contact angle decreases (Fig. 3c, brown), consistent with experiments (Fig. 1e,f), and the durotactic velocity increases (Fig. 3c, red). The speedup of durotaxis at low stiffness is thus favored by surface tension. Increasing it by increasing the pressure sensitivity to stiffness *s_P_* indeed yields smaller contact angles and hence faster durotaxis (Fig. 3e). The positive feedback between spreading and durotaxis produces a fast growth of the durotactic velocity, up to the regions where the surface tension contribution changes sign, at the contact angle of 90° (Fig. 3a, middle). After this point, surface tension points inwards and no longer promotes spreading (Fig. 3a, right); the spreading velocity therefore decreases (Fig. 3c, blue), and durotaxis slows down. Altogether, the combined effects of three-dimensional active wetting and the saturation of cellular forces at high stiffness explain why the durotactic velocity first increases and then decreases with substrate stiffness, as observed in our experiments (Fig. 2e,f), with a maximum around the contact angle of 90° (Fig. 3c).

The dependence of durotaxis on cluster size, cellular contractility, and stiffness gradient is also captured by our model. First, larger clusters have a larger active traction difference across them, which produces a higher durotactic velocity and shifts its maximum towards higher substrate stiffness (Fig. 3f), as in our experimental results (Fig. 2e). Second, the ROCK inhibitor Y-27632 reduces myosin-generated cellular contractility, which we implement by reducing the magnitude of all active forces in the model: monolayer contractility *ζ*, active traction *ζ_i_*, and tissue surface tension *γ*. Both contractility and active traction are active forces that would vanish completely with no myosin activity. In contrast, surface tension also has a passive contribution from cell-cell adhesion^53^, which produces a significant tissue surface tension even without myosin activity. Accordingly, we decrease contractility and active traction by a larger factor than surface tension. These parameter changes allow us to recover the decrease of durotactic velocity and the shift of its maximum towards lower stiffness (Fig. 3g) observed in our experimental results (Fig. 2f). Finally, increasing the stiffness gradient in the model produces faster durotaxis (Fig. 3h), which is consistent with our experimental measurements (Fig. 2g).

All in all, our model shows how the interplay between cell contractility, cluster size, stiffness, and cell traction forces can position clusters near contact angles of *θ* = 90°, and thus provide them with the sweet spot where cluster durotaxis is maximal.

### Cell clusters also durotax on ECM ligands close to neutral wetting

According to our theory, the emergence of fast durotaxis in the neutral wetting regime should be independent of the nature of the adhesion ligand. We thus tested whether a similar phenomenology can be observed on ECM ligands rather than on E-cadherin. Cell clusters were seeded on polyacrylamide gels of 0.2, 6, 24 and 200 kPa functionalized with fibronectin. Clusters displayed higher wettability on fibronectin than on E-cadherin substrates, fully wetting the substrates except for the lowest stiffness (Fig. 4a). To push the neutral wetting regime to an intermediate stiffness range, we treated clusters with 10 ng/mL of human Epidermal Growth Factor (hEGF), which increases cell contractility^54,55^ and cluster surface tension (Extended Data Fig. 6). With this treatment, cluster morphology showed a dependence with stiffness similar to that observed on E-cadherin substrates; full dewetting on soft substrates and a crossover to high wettability at intermediate stiffness (Fig. 4a,b). Cluster velocity was maximum in this intermediate regime, displaying a non-monotonic relationship with stiffness similar to that observed on E-cadherin substrates (Fig. 4c; Supplementary video 8). We then asked whether hEGF-treated clusters display durotaxis on substrates with a stiffness gradient coated with fibronectin. We found that this was indeed the case (Fig. 4d,e; Supplementary video 9). Like for cadherin-coated substrates, durotaxis was non-monotonic and peaked at the neutral wetting regime (Fig. 4f). Taken together, these data demonstrate that the durotaxis mode observed on E-cadherin gels is not a unique feature of cell-cell adhesion ligands. Rather, our data support that this behavior is a generic feature of clusters at the crossover between low and high wettability, whose location is tuned by substrate properties and active forces.

**Figure 4:**
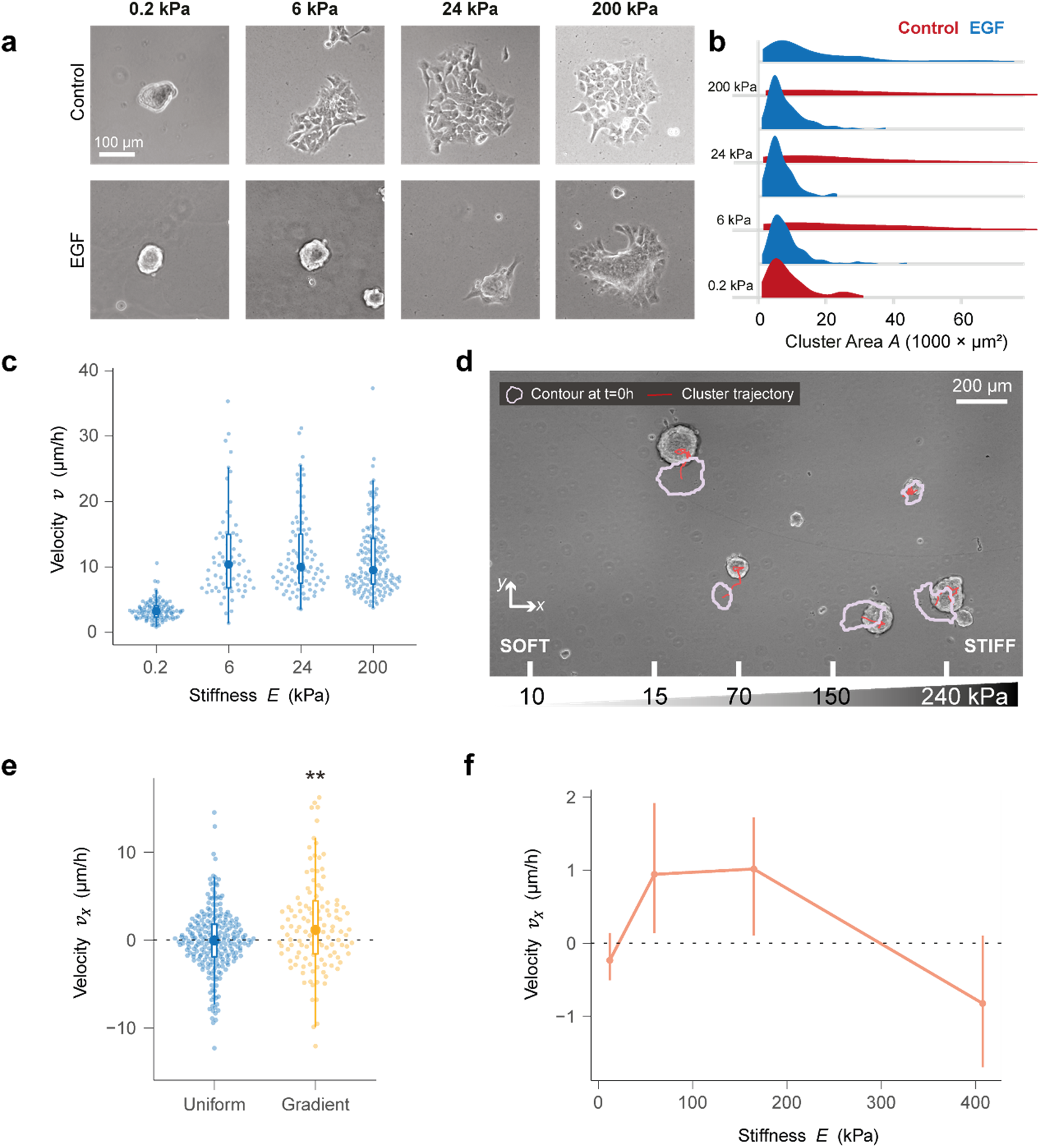
Wetting-based durotaxis also occurs on ECM substrates at high contractility. **a**, Representative phase-contrast images of A431 cell clusters seeded on fibronectin-coated uniform-stiffness gels of 0.2, 6, 24 and 200 kPa. **b**, Cluster area distribution for control and 10 ng/ml hEGF treatment. Clusters treated with hEGF spread less at lower stiffnesses than control clusters. **c**, Velocity of hEGF-treated A431 cell clusters on substrates of different stiffness. Velocity was computed at 100 min intervals. Each dot represents a cluster. For the sake of visualization, points above 99.5% percentile are not displayed. n=109 clusters (0.2 kPa), n=68 clusters (6 kPa), n= 89 clusters (24 kPa), n= 157 clusters (200 kPa). **d**, Representative image of hEGF-treated A431 cell clusters migrating on a fibronectin-coated substrate with a stiffness gradient. Image was taken at 10h. The original position (0h) of each cluster is represented by a purple outline. The red line represents the trajectory obtained by timelapse microscopy. Bottom scale indicates the local stiffness at each point of the substrate. **e**, Velocity along the *x*- axis of hEGF-treated A431 cell clusters, represented by dots, on uniform stiffness gels (pooled data from 0.2, 6, 24 and 200 kPa gels, n=245 clusters) and on stiffness gradients (n=128 clusters). Small dots represent individual clusters. The central dot in front of the boxplot represents the median. For the sake of visualization, points above 98% percentile are not displayed. (Permutation t-test (two tailed), p = 0.0027). **f**, Durotactic velocity as a function of stiffness. Data is median ± 95% CI estimated by bootstrapping; Bin 1 (0 < *E* ≤ 35 kPa): n=105 clusters (686 displacements). Bin 2 (35 < *E* ≤ 100 kPa): n=50 clusters (310 displacements). Bin 3 (100 < *E* ≤ 250 kPa): n=86 clusters (438 displacements). Bin 4 (*E* > 250 kPa): n=46 clusters (222 displacements).

### Sudden detachments of protrusions give rise to long durotactic hops

Finally, we studied the statistics of cell cluster movements. To this end, we computed the probability distribution of the displacements (*ρ*) of clusters on stiffness gradients coated with E-cadherin (Fig. 5). We found that most displacements were well described by an exponential distribution (Fig. 5a), as previously shown during random migration of single cells^56,57^. However, this distribution fails to capture long displacements (> 25 μm), which are much more frequent than expected. This fat tail in the distribution, which accounts for 0.4% of the total displacements, corresponds to hops that result from sudden retraction of cluster protrusions (Fig. 5b; Supplementary video 10). Strikingly, we found that these large hops are more durotactic than small displacements (Fig. 5c,d). Thus, cluster durotaxis arises from a combination of frequent small displacements and rare large hops.

**Figure 5.**
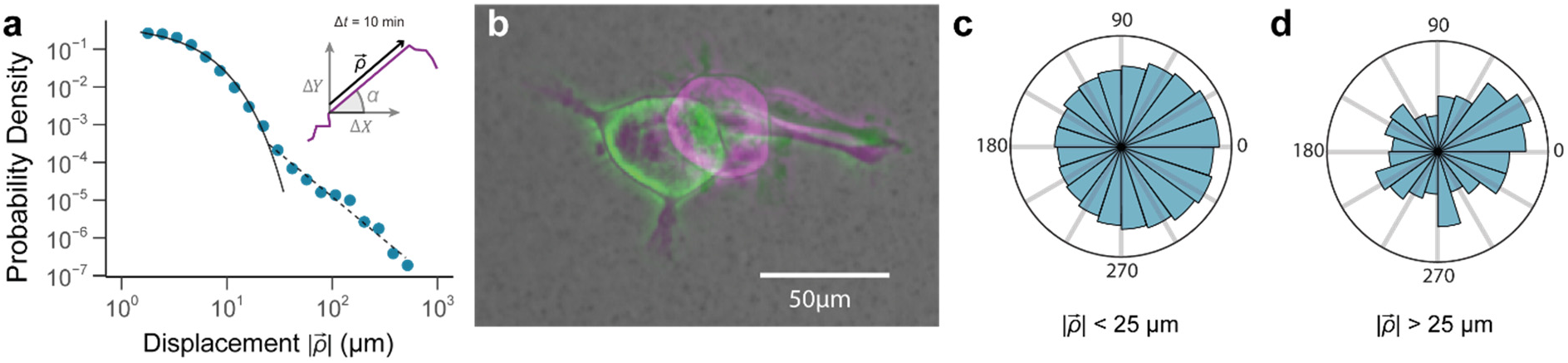
Sudden protrusion detachments give rise to long durotactic hops. **a**, Probability density of the displacements *ρ* (see inset) undergone by migrating cell clusters on stiffness gradient gels. Above displacements of 25 μm, the probability distribution deviates from an exponential (solid line) and is captured by a power-law function with exponent −2.31±0.14 (dashed line). Accordingly, we defined displacements *ρ* > 25 μm as long hops. **b**, Example of a sudden detachment of filopodia-like structures that results in a long hop. Green cluster is the initial image and the magenta one depicts the same cluster 10 min later. **c-d**, Distribution of the angle *α* between the instantaneous velocity vector and the *x*-axis (see inset of panel **a**) for short (**c**) and long hops (**d**).

## Discussion

Collective durotaxis is increasingly recognized as a key mechanism to drive directed cell migration in development^8,9^ and disease^1,11^. In this study, we provided a systematic analysis of collective durotaxis as a function of the physical properties of the cluster (size, three-dimensional shape, contractility) and of the substrate (stiffness, stiffness gradient, ligand identity). The analysis of hundreds of clusters revealed that durotaxis is a non-monotonic function of local substrate stiffness with a peak at intermediate stiffness. We showed that the position of the peak can be shifted to higher or lower stiffness by tuning cluster size and active forces. We developed a 3D model that captures the phenomenology in terms of the physics of active wetting.

As predicted in previous theoretical work^14,15^, our results experimentally demonstrate a mode of durotaxis in which cell clusters move as a whole, with the front and rear edges displacing in the same direction. This behavior is different from early studies of collective durotaxis using flat monolayers, which showed asymmetric spreading rather than directed migration^12,13^. Moreover, we generalize the theory of active wetting^31^ to 3D clusters including tissue surface tension to account for partial wettability, i.e. finite contact angles. We show that the interplay between wettability and substrate stiffness is essential to explain the non-monotonic behavior of durotaxis. This wetting-based mode of durotaxis may involve but does not require long-range force transmission across the tissue. This feature distinguishes it from previous models of durotaxis that imposed global force transmission by treating the cell cluster as an elastic medium^12,16,58–60^. Overall, our work provides a theory for stiffness-dependent 3D active wetting that explains the measured properties of collective cell durotaxis, and may serve as a guide for further experiments. These experiments should seek to understand, for example, whether the width of the peak can be tuned as predicted by the model, which we were unable to analyze in this work due to high scattering of the data. In the future, the picture reported here can be extended to account for flows resulting from spatial pressure variations across a cluster, for the size-dependent surface tension of cell aggregates^61^, and for elastocapillary effects resulting from substrate deformation^62–64^.

Our study demonstrates that durotaxis can be mediated not only by ECM ligands but also by E-cadherin. E-cadherin migration is relevant in early developmental stages in which the ECM is absent or weakly expressed. The paradigmatic example of collective cell migration through cadherin receptors is border cell migration during *Drosophila* oogenesis^25,26^. In this process, the cohesive border cell cluster migrates antero-posteriorly in the egg chamber by establishing dynamic protrusions with neighboring nurse cells through E-cadherin receptors. In zebrafish, E-cadherin also mediates the migration of progenitor cells and of cell sheets during epiboly^65,66^. In cancers that retain epithelial features, E-cadherin is the dominant adhesion molecule between cancer cells and is therefore responsible for local and global migratory events and rearrangements within the tumor^30,67^. Besides E-cadherin, N-cadherin has also been shown to mediate long distance migration of inter-neuron precursors and neurite outgrowth^29,68,69^. In all these processes, the mechanisms driving directed cell migration remain incompletely understood. Whereas determining the presence of mechanical gradients *in vivo* presents outstanding experimental challenges, our results establish durotaxis as a plausible mechanism underlying directed cell migration based on cadherin adhesion.

We studied whether the existence of optimal cluster durotaxis in the neutral wetting regime is independent of the nature of the adhesion ligand by carrying out experiments on substrates coated with E-cadherin and fibronectin. Clusters on E-cadherin exhibited full dewetting on all substrates except for the stiffest ones. By contrast, clusters on fibronectin fully wetted all substrates except for the softest ones. Several molecular mechanisms could explain these differences. These include differences in coating density, in affinity of cadherin-cadherin vs fibronectin-integrin bonds^70^, in adhesion reinforcement mechanisms^71^ (e.g. talin vs alpha-catenin unfolding), in lateral adhesion clustering^72^ or in adhesion regulation by growth factors^73^. Irrespective of the molecular mechanisms at play, an increase in contractility with hEGF brought clusters on fibronectin-coated substrates to wettability conditions similar to those on cadherin-coated substrates, showing full dewetting on soft substrates, full wetting on stiff ones, and neutral wetting at intermediate stiffness. Under these conditions, the non-monotonic behavior of durotaxis was recovered, indicating that this behavior is independent of the nature of the adhesion ligand.

Inspection of the distribution of cluster displacements showed that collective durotaxis involves a combination of small but frequent displacements following an exponential distribution and large but rare hops following a power law. Dynamics featuring rare and sudden reconfiguration events have been described in disordered systems including rearrangements in colloidal glasses, avalanches in sandpiles, and earthquakes in geological systems^74,75^. These apparently distinct materials, which span several orders of magnitude in size, share that they all respond to slow shear through some kind of stick-slip dynamics^76^. The data presented here suggest that these dynamics also take place in wetting processes in tissues, enabling the emergence of fast directed migration.

Our finding of an optimal regime for collective durotaxis provides a new strategy for the regulation of directed cell migration *in vivo*. By tuning the local stiffness of a substrate or the active properties of a cluster, organisms can finely control the onset of directed migration and its extent^77^. Conversely, abnormal tissue stiffening or softening, or changes in active cluster mechanics could impair physiological migration or trigger undesired durotaxis. A particularly relevant pathological context for our findings concerns the movement of cellular clusters during cancer invasion and metastasis. Our results indicate that these clusters will exhibit poor migration when fully wetting or dewetting their microenvironment, but will be able to follow stiffness gradients efficiently at the crossover between low and high wettability. Further work is needed to study the wettability of cellular clusters *in vivo* and the implications in physiological and pathological directed cell migration. Our study provides a general physical framework to address this question.

## Acknowledgments

We thank all the members of our groups for their discussions and support. We thank Anghara Menéndez, Susana Usieto and Beatriz Martin for technical assistance and Anabel-Lise Le Roux for helping to produce and purify histidine-tagged mCherry. We also thank Erik Sahai for sharing cell lines and plasmids used in this work. Finally, we thank Juan Francisco Abenza, Eleni Dalaka and Tom Golde for their feedback on the manuscript. This paper was funded by the Generalitat de Catalunya (AGAUR SGR-2017-01602 to X.T., AGAUR SGR-2017-1061 to J.C., the CERCA Programme, and “ICREA Academia” awards to P.R-C. and J.C.); Spanish Ministry for Science and Innovation MICCINN/FEDER (PGC2018-099645-B-I00 to X.T., PID2019-110298GB-I00 to P.R-C., PID2019- 108842GB-C21 to J.C., RTI2018-101256-J-I00 and RYC2019-026721-I to R.S., FPU19/05492 to I.P-J., FPU15/06516 to M-E.P.); Fondo Social de la DGA (grupos DGA) to V.G. and J.M.F.; European Research Council (Adv-883739 to X.T.); Fundació la Marató de TV3 (project 201903-30-31-32 to X.T.); European Commission (H2020-FETPROACT-01-2016-731957 to P.R-C. and X.T.); the European Union’s Horizon 2020 research and innovation programme (under the Marie Skłodowska-Curie grant agreement no. 797621 to M.G-G.); La Caixa Foundation (LCF/PR/HR20/52400004 to to P.R-C. and X.T.); IBEC is recipient of a Severo Ochoa Award of Excellence from the MINECO. R.S. is a Serra Húnter fellow.

## Author contributions

M-E.P., R.S. and X.T. conceived the project. M-E.P., R.S., and I.C.F. performed experiments. V.G., J.M.F. and P.R-C. contributed technical expertise, materials and discussion. I.P-J., R.A. and J.C. developed the model. M.G-G. and R.S. developed analysis software. M-E.P., I.P-J., R.A., R.S., J.C., and X.T. wrote the manuscript. All authors revised the completed manuscript. R.A., R.S., J.C. and X.T. supervised the project.

## Code availability

Analysis procedures and code implementing the model are available from the corresponding authors on reasonable request.

## Data availability

The data that support the findings of this study are available from the corresponding authors on reasonable request.

**Extended Data** is available for this paper

**Correspondence and requests for materials** should be addressed to R.A., R.S., J.C. or X.T.

## Methods

### Cell culture

A431 cells were cultured in Dulbecco’s Modified Eagle’s Medium containing high glucose and pyruvate (11995, Thermofisher) supplemented with 10% fetal bovine serum, 100 units·ml^-1^ penicillin and 100 μg·ml^-1^ streptomycin. Cells were maintained at 37°C in a humidified atmosphere containing 5% CO_2_. Prior to experiments, cells were starved for 24 hours in starvation media (Dulbecco’s Modified Eagle’s Medium containing high glucose and pyruvate supplemented with 1% FBS, 100 units·ml^-1^ penicillin and 100 μg·ml^-1^ streptomycin). Versene (15040066, Gibco) was used to collect cells from flasks as a non-enzymatic cell dissociation reagent aiming to preserve the integrity of membranal E-cadherin molecules prior to cell seeding on gels functionalized with E-cadherin.

### Cell clusters formation

A431 heterogeneous cell clusters were obtained by seeding 5·10^3^ cells/well in Corning Costar Ultra-Low Attachment Multiple Well Plate (CLS3474-24EA) in starvation media. Cell clusters were mechanically disaggregated into smaller clusters exhibiting heterogeneous sizes by pipetting up and down with a series of pipette tips of different sizes. Cellular debris was discarded by centrifuging disaggregated clusters at 0.3 rpm for 0.5 min.

### Cell clusters seeding

Cell clusters were resuspended in media containing 5 μM RHO/ROCK pathway inhibitor (Y-27362) and seeded on stiffness gradients in a total volume of 50 μL to allow for cluster adhesion to the entire surface of the gradient, thus covering the whole range of stiffness. 45 minutes later, 1 mL of media containing 5μM RHO/ROCK pathway inhibitor (Y-27362) was added to prevent the gels from drying out. After 1 hour, Y-27362 was carefully removed by slow aspiration and 1.5mL of fresh starvation medium was added. Clusters were imaged 2 hours later.

### Lentiviral transfection for stable Lifeact-mCherry expression

HEK293T cells were transfected as previously described^78^ to produce lentiviral particles inducing stable expression of Lifeact-mCherry. A431 wild type cells were infected as previously described^78^. Two weeks later, infected cells were sorted using an ARIA fluorescence-activated cell sorter (BD) aiming to select those cells exhibiting similar fluorescence intensity.

### Cell adhesion assay

A431 cells were resuspended at a concentration of 10^6^ cells/mL and incubated for 15min in ice-cold starvation media containing either 40 μg/mL α-GFP antibody (A10262, Life technologies; Control) or 40 μg/mL DECMA-1 antibody (U3254, Sigma Aldrich). Cells incubated with control or DECMA-1 antibody were seeded on gels functionalized with E-cadherin. Cell adhesion was allowed for 15min before gently washing three times with 1xPBS containing calcium and magnesium, and the remaining adhered cells were fixed with 4% PFA in 1xPBS containing calcium and magnesium, followed by Hoecsht staining for nuclei quantification. An inverted microscope (Nikon Eclipse Ti) equipped with a 2x 0.06NA objective was used to image the entire gel surface. An intensity threshold was set on FiJi software to binarize the images and automatically count the number of adhered cells per condition. Cell density was assessed, data sets were transformed to obtain normal distributions and non-parametric statistical tests were performed.

### Glass-bottom dish silanization

Glass-bottom dishes (P35-0-20, Mattek) were silanized using a 2:1:80 solution of acetic acid/bind-silane (M6514, Sigma)/ethanol for 30 min. The dishes were washed twice with ethanol and dried by aspiration.

### Polyacrylamide gels of uniform stiffness

A 1 mL gel premix solution containing 2% bis-acrylamide and 40% acrylamide (proportions vary according to desired stiffness; see Table 1), 15μL irgacure 5% w/v (BASF, Germany), 6 μL acrylic acid (147230, Sigma Aldrich), 84 μL 1M NaOH and 10 μL 500nm-diameter yellow-green fluorospheres was prepared (F8813, Thermofisher). A drop of 16 μL of gel premix was added to the center of the previously silanized glass-bottom dish, and an 18-mm diameter glass coverslip treated with Repel Silane (General Electric, USA) was placed on top to distribute the volume evenly and flatten the gel surface. Glass-bottom dishes containing a sandwich of gel premix were placed under UV light for 5min to allow for gel polymerization. Next, 10x PBS was added and round-tip tweezers were used to separate top coverslips from the gels polymerized on glass-bottom dishes.

**Table 1.**
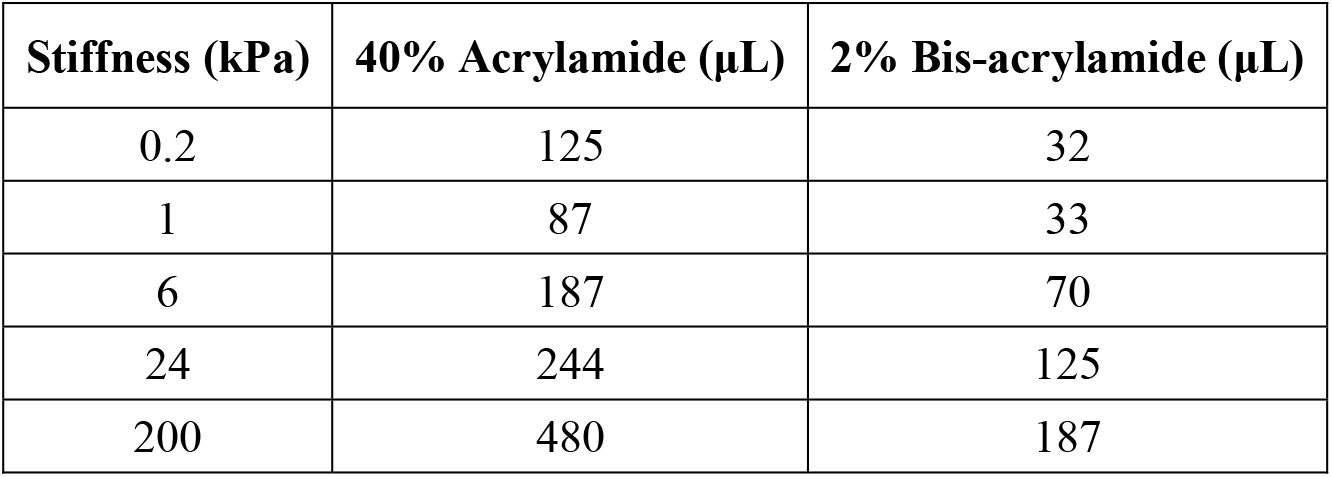
Stiffness and corresponding Acrylamide/Bis-acrylamide concentrations for 1mL gel premix.

### Polyacrylamide gels with a stiffness gradient

A 25μL drop of gel premix containing 15% acrylamide, 1% bis-acrylamide, 0.75 mg·mL^-1^ irgacure, 0.60% acrylic acid, 100mM NaOH and a dilution of 1:100 from stock 500nm-diameter fluorescent beads (F8813, Thermofisher) was added to the center of previously silanized glass-bottom dishes and covered with an 18mm-diameter glass coverslip treated with Repel Silane (General Electric, USA). Gradients of stiffness on polyacrylamide gels were obtained as previously described^12,43^. Briefly, making use of an opaque sliding mask during UV-triggered gel polymerization we polymerized gels exhibiting a gradient of stiffness. To obtain “shallow” and “steep” gradients, the opaque mask was moved at rates of 50 μm·s^-1^ and 30 μm·s^-1^, respectively. Finally, 10xPBS was added to facilitate the removal of top coverslips using round-tip tweezers. Gel stiffness was measured with AFM after every experiment.

### Gel functionalization with oriented E-cadherin

To functionalize uniform and stiffness gradient gels we used a previously described method involving carbodiimide reactions^79^. Briefly, a mix of 20 mM Nα,Nα-Bis(carboxymethyl)-L-lysine hydrate (NTA-NH2, 14580, Sigma-Aldrich) and 25 mM Copper (II) Sulphate 5-hydrate (CuSO_4_, 131270, Panreac) in 1xPBS buffer was brought to pH=10.0 and centrifuged at 4500 rpm for 15min. The pellet was discarded while the supernatant (formed by a solution containing NTA-NH_2_/Cu^2+^ complex) was brought to pH=7 and filtered using a 0.20 μm diameter filter. In parallel, previously polymerized polyacrylamide gels were incubated with 100mM N-(3-dimethylaminopropyl)-N’-ethylcarbodiimide hydrochloride crystalline, imidazole (EDC, E6383, Sigma-Aldrich) and 200mM N-Hydroxysuccinimide (NHS, 130672, Sigma-Aldrich) in 20mM Hepes pH=7.0 buffer at 37°C for 20min. Then, the gels were quickly washed twice with 1xPBS and incubated at 37°C with filtered NTA-NH_2_/Cu^2+^ solution aiming to bind it covalently to carboxyl groups on the gel surface. Two washes with 1xPBS were performed 45min later, followed by addition of 1M Tris(hydroxymethyl)aminomethane hydrochloride (TRIS, 648317 Merck) pH=8.0 for 30min to hydrolyze unreacted active carboxyl groups. Next, the gels were washed twice with 1xPBS and incubated at room temperature with a drop of 25μL of 0.01mg·mL^-1^ histidine-tagged E-cadherin (8505-EC-050, R&D Systems) covered with rectangles of parafilm, ensuring an even distribution of the drop across the entire surface of the gel. E-cadherin oriented chelation on the gel was achieved after 1 hour of incubation, and excess E-cadherin molecules were rinsed with two washes of 1xPBS. The metal chelation of its C-terminal polyhistidine tail, via the metal chelator complex (NTA- Cu^2+^) previously introduced at the polyacrylamide gels surface, guarantees the proper orientation of the protein fragment by mimicking its presentation on cell surfaces. However, the polyhistidine-NTA interaction is highly dependent on pH, ionic strength, and other media conditions. Thus, the gels were incubated with 50mM EDC and 75mM NHS in 20mM pH=7.0 Hepes buffer at 37°C for 45 min aiming to covalently bind E-cadherin molecules to the gels by promoting amide link formation between amino groups in the polihistidine tag and the carboxylated groups presented in the NTA. Next, the gels were washed twice with 1xPBS and incubated with 1M imidazole/1mM EDTA in PBS for 20min to compete for/chelate copper ions and thus rinse non-covalently bound E-cadherin molecules, followed by two 1xPBS washes. Finally, the gels were passivated with 0.1 mg·mL^-1^ pLL-g-PEG, sterilized with UV and used for experimental purposes within the 24h after the functionalization process.

### Gel functionalization with fibronectin

Polyacrylamide gels were functionalized using carbodiimide reactions. Briefly, gels were incubated with 100 mM EDC and 200 mM NHS in 20 mM Hepes pH=7.0 buffer at 37°C for 20 min. Next, gels were quickly washed twice with 1xPBS and incubated at 37°C with a dilution of 0.1 mg/mL fibronectin (33016015) in 1xPBS for 45 min. Finally, gels were washed twice with 1xPBS and incubated with 1M Tris pH=8.0 for 30 min at RT, followed by two 1xPBS washes. Gels were kept overnight at 4°C prior to UV-sterilization and cluster seeding.

### Histidine-tagged GFP and histidine-tagged mCherry production and purification

Histidine-tagged GFP and histidine-tagged mCherry were produced in *Rosetta E. coli* strain and purified using Ni-NTA columns, as previously described^80^.

### Protein incorporation quantification in polyacrylamide gels

To study the protocol’s functionalization efficiency, histidine-tagged E-cadherin was replaced by Histidine-tagged GFP, whose functionalization and orientation are achieved likewise, offering a direct fluorescence readout for the protocol validation. Aiming to provide a realistic idea of the extent of protein incorporation, their molar concentrations were normalized. Fluorescence images were taken from functionalized gels using an inverted microscope (Nikon Eclipse Ti) equipped with a 10x 0.30 NA objective (for stiffness gradient gels) or a 20x 0.45 NA objective (for uniform stiffness gels).

Further experiments were carried out to assess the functionalization protocol in a stepwise manner. Polyacrylamide gels depleted of acrylic acid were not functionalized with histidine-tagged GFP molecules, indicated by a decay in fluorescence intensity. A second round of EDC/NHS activation was carried out aiming at the formation of a covalent bond between histidine-tagged GFP molecules and the gel. The decay in fluorescence intensity after imidazole-mediated rinse in gels that did not undergo a second round of EDC/NHS activation suggested that histidine-tagged GFP orientation was achieved as a result of histidine chelation of histidine-tagged GFP molecules by NTA/Cu^2+^ complexes. Maximum fluorescence intensity values were obtained after a second round of EDC/NHS activation in gels that were not rinsed with imidazole/EDTA. A mild decay of fluorescence intensity experienced by an imidazole/EDTA-mediated rinse in gels that underwent a second round of EDC/NHS activation suggested that a significant part of histidine-tagged GFP molecules were successfully oriented and covalently bound to the gels after the second round of EDC/NHS activation (Extended Data Fig. 2b).

### Contractility enhancement and inhibition experiments

To study the effect of contractility on cluster migration, 0.5 μM Y-27632 ROCK inhibitor and 1.65 nM human epidermal growth factor (hEGF) were used to inhibit and enhance cell contractility, respectively.

### Time lapse microscopy

Multidimensional acquisition routines were performed on automated inverted microscopes (Nikon Eclipse Ti) equipped with thermal, CO_2_ and humidity control using MetaMorph, Micromanager and NIS-Elements softwares. Time-lapse experiments started approximately 4h after cell seeding. The image acquisition interval was set to 10 min, and a typical experiment was run for at least 14h. Images were acquired using a 10x 0.3 NA objective, and an automated stage was used to save 3 overlapping stage positions that covered ~3 mm of the stiffness gradients starting from the soft edge.

### High-resolution images of cell clusters

An inverted Nikon microscope equipped with a spinning disk confocal box (CSU-WD, Yokogawa) was used to acquire high resolution images of A431 mCherry-Lifeact cell clusters seeded at different stiffness. A z-step equal to 0.2 μm was acquired for every cluster, ensuring to capture whole cell clusters.

### Contact angle measurement

To compute the contact angle (*θ*) formed between the substrate and the line tangent to the edge of the cluster (Fig. 1e, inset) we acquired high resolution stacks of mCherry-Lifeact A431 cell clusters seeded on 0.2, 1, 6, 24 and 200 kPa (the *z*-step was 0.26 μm after correcting for the focal plane shift). Extended Data Fig. 7 shows the basic strategy used to calculate the contact angle as a function of the cluster radius (*R*_sphere_), the contact radius (*R*) and the cluster height (*H*). In dewet clusters, we estimated *R*_sphere_ from the maximum Z projection of all the images in the stack and *R* from the basal plane (Extended Data Fig. 7a). Then, the contact angle was given by 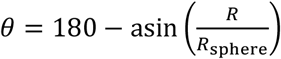. For wet clusters, we estimated *R*_sphere_ from the height *H* and *R* (Extended Data Fig. 7b). Both *H* and *R* were measured from the Z stack. Then, the contact angle was given by 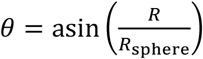.

### Traction force microscopy

Traction forces were computed using Fourier transform based traction microscopy with a finite gel thickness as previously described^81^. Gel displacements experienced between any experimental timepoint and a reference image of the relaxed state of beads after cell cluster trypsinization were computed using home-made particle imaging velocimetry software^81^.

### Stiffness gradient profile measurement with Atomic Force Microscopy

Stiffness gradients and uniform stiffness gels were mapped individually using a commercial Atomic Force Microscope (AFM) (JPK Nanowizard) operated as previously described^82,83^. Briefly, a V-shaped cantilever (Bruker) with a triangular tip and a spring constant of *k* = 0.03 N·m^-1^ was used to indent the gels to ensure the cantilever deflection resulted within the linear detection range of the AFM. The cantilever spring constant was calibrated using a thermal fluctuation method. The relationship between the photodiode signal and cantilever deflection was computed from the slope of the force displacement curve obtained at a region without gel sample. For each sample, 5 force-displacement (F-z) curves (where *F*= *k*· *d*, d being the deflection and *z* being the piezotranslator position) were acquired by ramping the cantilever forward and backward at a constant speed (5 μm amplitude, 1 Hz and approximately 1 μm of indentation). Each experimental *F-z* curve was fitted to the four-sided pyramidal indenter model:

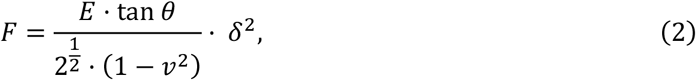

where *E* is the Young’s modulus, *v* is the Poisson’s ratio, *θ* is the semi-included angle of the pyramidal indenter and *δ* is the indentation depth. The parameter *v* is assumed to be 0.5, and the indentation depth is calculated as *δ* = *z* – *z*_0_ – *d*, where *z*_0_ is the tip-gel contact point. *E* and *z*_0_ were estimated by leastsquares fitting of this equation to the F-z curve recorded on each gel point. Young’s modulus was measured every 200 μm along the axis of maximum gel stiffness change.

### Cluster tracking using phase images

Custom-made Matlab scripts combined with “Grid/Collection stitching” plug-in from FiJi software were used in order to register and stitch three overlapping images covering 3 mm of the stiffness gradients. Briefly, time-lapse images from the green fluorescent channel containing fluorescent beads (F8812, ThermoFisher) were registered using a custom-made Matlab script. “Grid/Collection stitching” plugin from FiJi was used to stitch registered maximum intensity images from fluorescent beads, and a custom-made Matlab script was run in order to stitch phase contrast images using the *XY* coordinates provided by FiJi.

Stitched phase contrast images were used to segment clusters. Images were treated with gaussian and standard deviation filters to find cluster edges. After applying automatic thresholding and detection algorithms we detected cluster position. Clusters were linked based on proximity, and tracks were generated and labelled with sequential numbers. Segmented images with cluster labels were merged with phase contrast images, and clusters were manually selected using the label number. Inconsistently tracked and incorrectly segmented clusters were discarded for analysis. Clusters whose tracks left the field of view were kept until the timepoint in which their outline overlapped with the image boundary, whereas clusters whose tracks interacted were kept until the timepoint in which the interaction took place. Final tracks contained the *x* and *y* position for each cluster at the measured timepoints. Noise in trajectories was estimated at 1.5 μm by tracking pieces of immobile debris.

### Image processing of high-resolution images of clusters

Acquired images of mCherry-Lifeact A431 cell clusters were processed using Imaris software. A gaussian filter was applied to the images to smoothen the fluorescence signal before generating a surface to visualize the tridimensional shape of clusters.

### Statistics

Statistical analyses and plotting were performed using R v 4.1.2 (R Core Team). For comparing two groups, we performed two-sample permutation t-test implemented in the “MKinfer” library. We assessed differences among groups using permutation-based analysis of variance (Permutation-based ANOVA), which does not require normality or homogeneity of variances, using the “lmperm” library. We ran a post hoc permutational test to assess the pairwise significance among groups, using the library “rcompanion” and reporting the adjusted p-value with the “false discovery rate” method. 95% confidence intervals of medians were estimated using bootstrap intervals of 10,000 resamples.

### Modelling dynamic evolutions

We assume that a spherical-cap cluster evolves while keeping the total volume fixed. The stiffness profile is chosen to be linear with position on the gel (Eq. (S15)) with its values inferred from experiments (Fig. 2a as a representative example). Traction, friction, and surface tension functions with the stiffness are chosen, linear (for simplicity, following Eqs. (S16)-(S18)) or saturated (Eq. (S12)). At each step, the velocity profile is computed solving Eq. (S6), with the polarity profile Eq. (S7) and the boundary conditions Eq. (S11). By a simple Euler integration algorithm with a time step Δ*t*, the center-of-mass *X* and contact radius *R* are evolved following

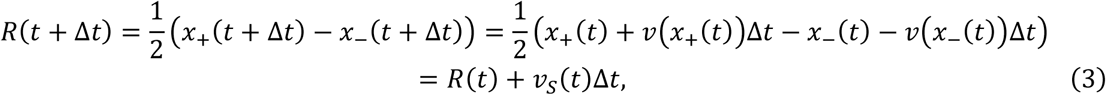

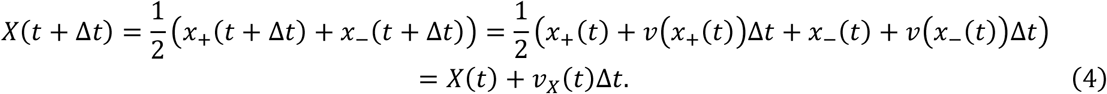

Imposing volume conservation with 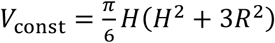, this fixes the new height, and then the contact angle is

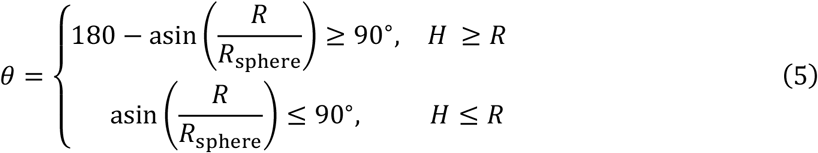

In general, the clusters do not strictly follow a quasistatic evolution, defined as a sequence of stages with vanishing spreading velocity (see for instance *v_S_* in Fig. 3c and Figs. S9-S11).

### Model parameters fit

We use the experimental measures from Table 2 as references to infer the dependencies of the active traction parameter *ζ_i_* and pressure *P* with stiffness, since *ζ_i_* ~ *T_r_/h* (where *h* is the height of the basal monolayer, typically *h* = 5 μm^31,81^) and *P* ~ *T_z_*. Through Laplace’s law, surface tension is 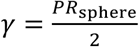 and so it will depend on the size of the clusters. From here, the slopes of these profiles might be slightly changed to explore a wider range of the phenomenology from the model. Typically, we take 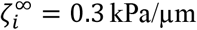^31,44^, a range of *E*\* = (50 – 450) kPa for the active traction saturation profile (Eq. (S12)), and a range of pressure sensitivity to stiffness of *s_P_* = (0.3 – 9.1) × 10^-2^ for the linear pressure increase (Eq. (S13)).

**Table 2.** Approximate experimental measures of the radial ***T_r_*** and vertical ***T_z_*** components of traction forces in cell clusters on top of uniform-stiffness substrates of 1 and 6 kPa (from Fig. 1i,j). Also, surface tension estimates for clusters of an apparent size of ***R_sphere_*** ~ **30** μm (Fig. 1g,h).

| Stiffness (kPa) | Traction $T_r$ (Pa) | Traction $T_z$ (Pa) | Surface tension $\gamma$ (mN/m) |
| --- | --- | --- | --- |
| 1 | 10 | 10 | 0.15 |
| 6 | 40 | 50 | 0.75 |

## Extended Data

### Extended data figures

**Extended Data Figure 1.**
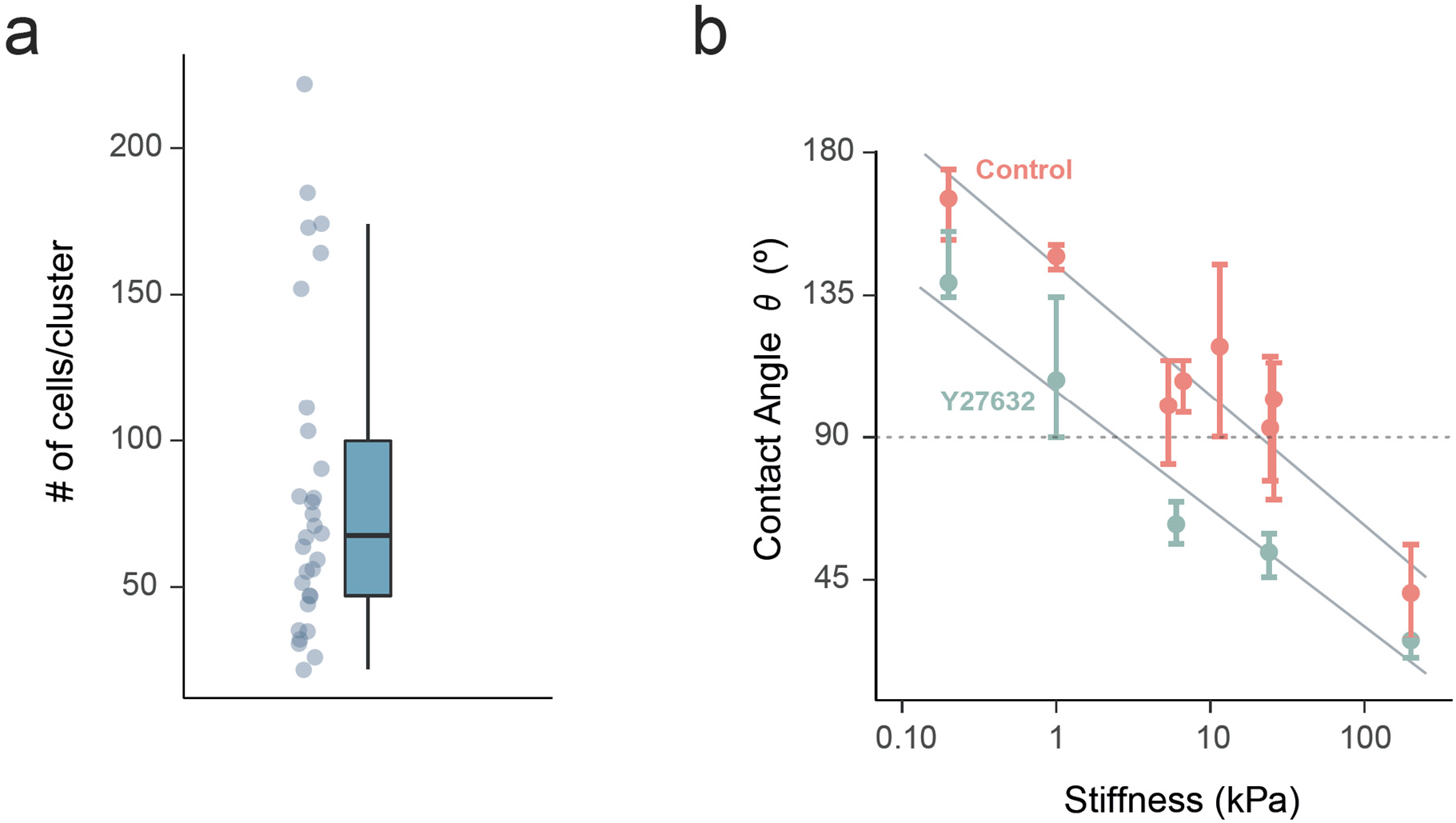
Characterization of cluster size and contact angle. **a,** Distribution of the number of cells per cluster (n=30 clusters). The high variability was intended in order to study the effect of cluster size on durotaxis. **b,** Contact angle as a function of stiffness for control cells (red) and for cells treated with 0.5 μg/ml of Y27632 (turquoise). Data are median ± 95% CI estimated by bootstrapping (n=4-9 clusters for control and n=8-34 clusters for Y27632).

**Extended Data Figure 2.**
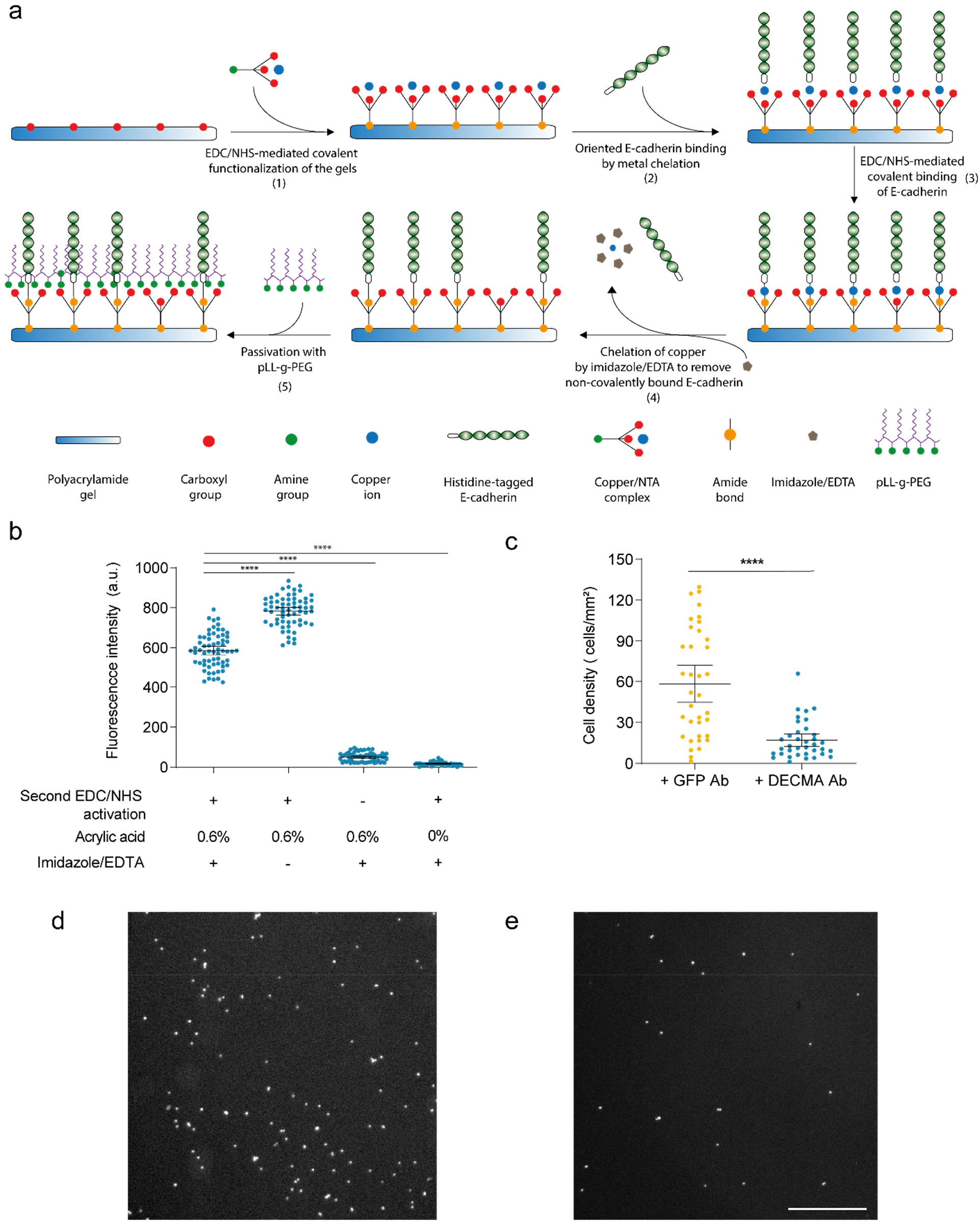
Functionalization of polyacrylamide gels with oriented E-cadherin extracellular domains (EC1-5). **a**, Scheme showing the protocol to covalently attach E-cadherin extracellular domains on the surface of a polyacrylamide gel mimicking its oriented presentation on cell surfaces. Briefly, polyacrylamide gels containing acrylic acid and thus presenting free carboxyl groups were activated with EDC/NHS and incubated with a solution of NTA/Cu ^2+^ complexes (1) aiming to form a covalent amide bond between carboxyl groups in the gels and amino groups from NTA/Cu ^2+^ complexes. Next, gels were incubated with histidine-tagged E- cadherin extracellular domains EC1-5 (2), which spontaneously oriented along NTA/Cu ^2+^ complexes through metal chelation of their poly-histidine tag. A covalent amide bond between histidine-tagged E-cadherin extracellular domains and NTA/Cu ^2+^ complexes was formed upon a second round of EDC/NHS activation (3). Finally, a solution containing imidazole/EDTA was used to rinse/elute non-covalently bound histidine-tagged E-cadherin extracellular domains (4) prior to gel passivation with pLL-g-PEG (5). **b**, Fluorescence intensity as a readout of protein incorporation in polyacrylamide gels including (+) or omitting (−) steps in the protocol. First row indicates whether gels underwent a second EDC/NHS treatment to covalently bind histidine-tagged GFP to the gels; second row indicates concentration of acrylic acid; third row indicates whether gels were rinsed with imidazole/EDTA to remove non-covalently bound histidine-tagged GFP. **c**, Adhesion assay performed on A431 cells in the presence of E-Cadherin blocking antibody (DECMA). This assay validates the specificity of our coating. For panels b-c, data is mean ± 95% CI. ****, p-value < 0.0001, Permutation test. **d-e**, Fluorescence image of nuclei (labelled with Hoechst) of attached A431 single cells in controls and DECMA-treated A431 single cells, respectively. Scale bar, 500 μm.

**Extended Data Figure 3.**
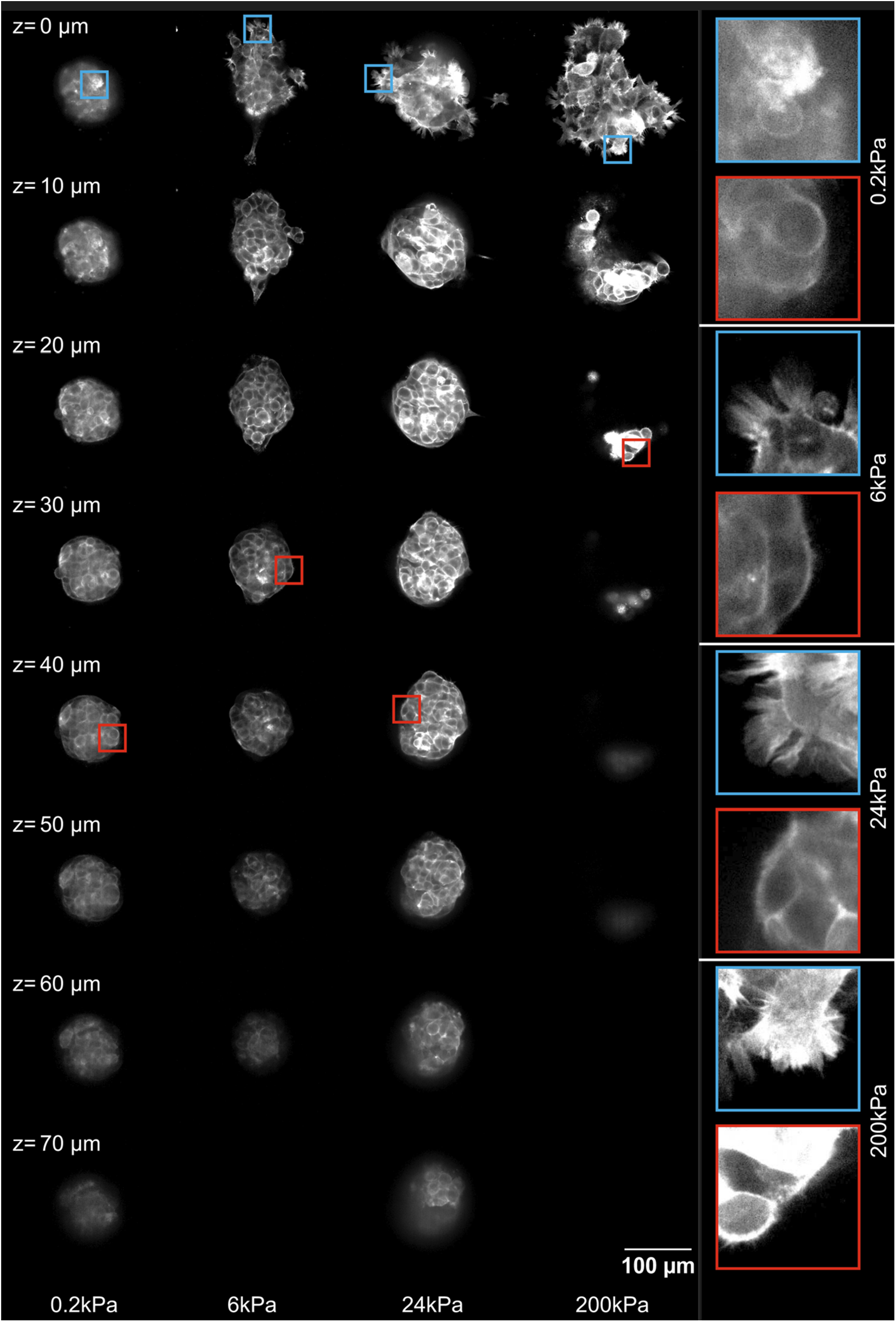
3D cluster profile. Z-stack of A431 clusters expressing LifeAct-mCherry for 0.2, 6, 24 and 200 kPa uniform stiffness gels coated with oriented E-cadherin. Slices are shown with a z-step size of 10 μm. Basal plane is z = 0 μm. Insets indicate zoomed areas of different planes.

**Extended Data Figure 4.**
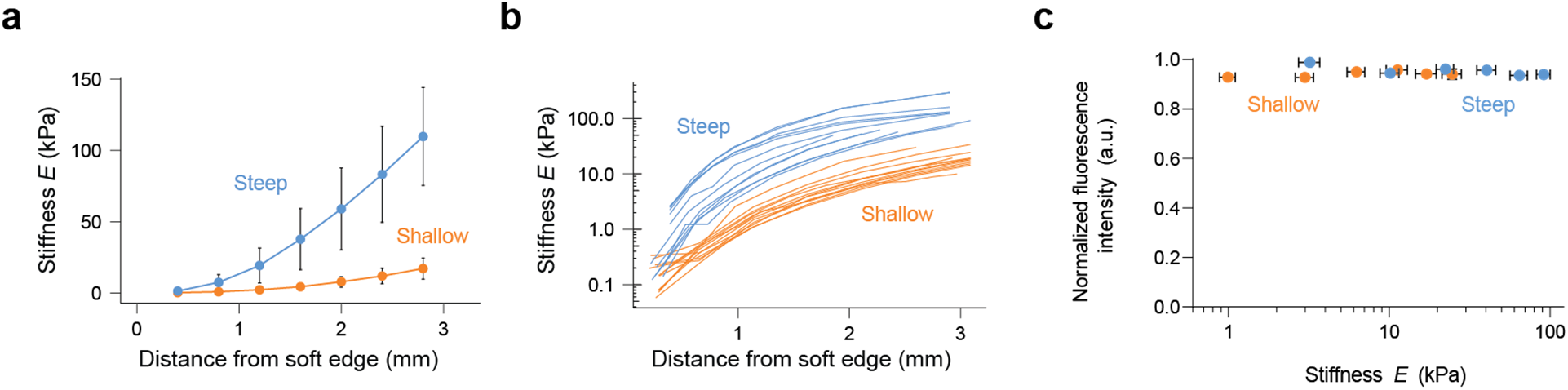
Stiffness profile and protein incorporation of shallow and steep gradient gels. **a**, Stiffness profile as a function of distance from soft edge for shallow (n=12, orange) and steep (n=12, blue) stiffness profiles. The stiffness profile was determined using AFM for every gel (see Methods). Data is mean ± SD. **b**, Stiffness profile for individual gels (shallow in orange, steep in blue). Note the logarithmic scale on the Stiffness axis for the sake of visualization. **c**, Normalized fluorescence intensity of histidine-tagged mCherry signal as a readout of protein incorporation as a function of stiffness for shallow and steep gradient gels.

**Extended Data Figure 5.**
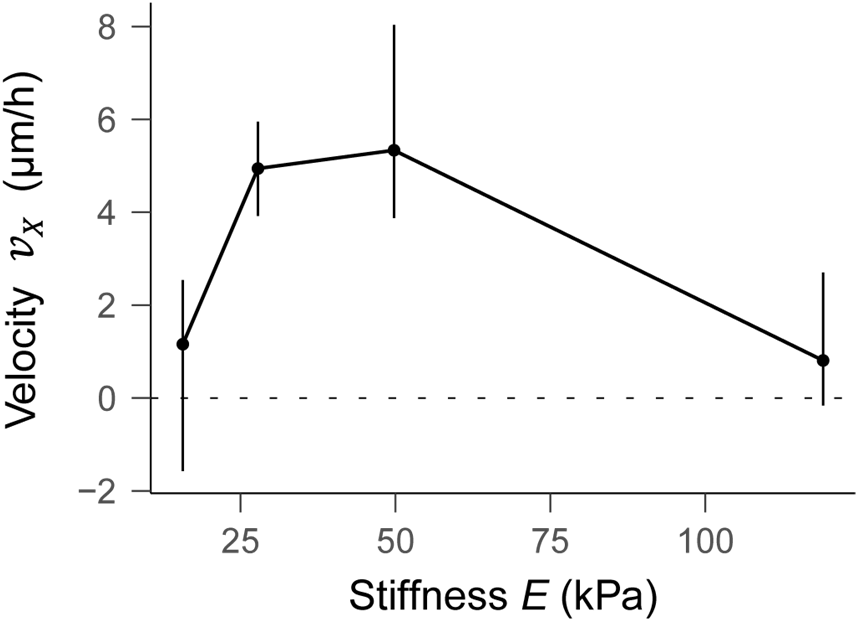
Dependence of durotactic velocity on stiffness for clusters within the 40 ± 10 kPa/mm gradient range. Data analyzed include only the positions of the gels where the gradient was 40 ± 10 kPa/mm, showing that the sharp increase of durotactic velocity with stiffness cannot be accounted for by changes in the gradient. Bin 1 (0 < *E* ≤ 20 kPa): n= 28 clusters (136 displacements), *E’* = 36.8 kPa/mm. Bin 2 (20 < *E* ≤ 40 kPa): n= 62 clusters (386 displacements), *E’* = 40.5 kPa/mm. Bin 3 (40 < *E* ≤ 80 kPa): n= 23 clusters (102 displacements), *E’* = 47.8 kPa/mm. Bin 4 (*E* > 80 kPa): n= 43 clusters (313 displacements), *E’* = 48.8 kPa/mm. Data includes clusters of all sizes. Data is median ± 95% CI estimated by bootstrapping.

**Extended Data Figure 6.**
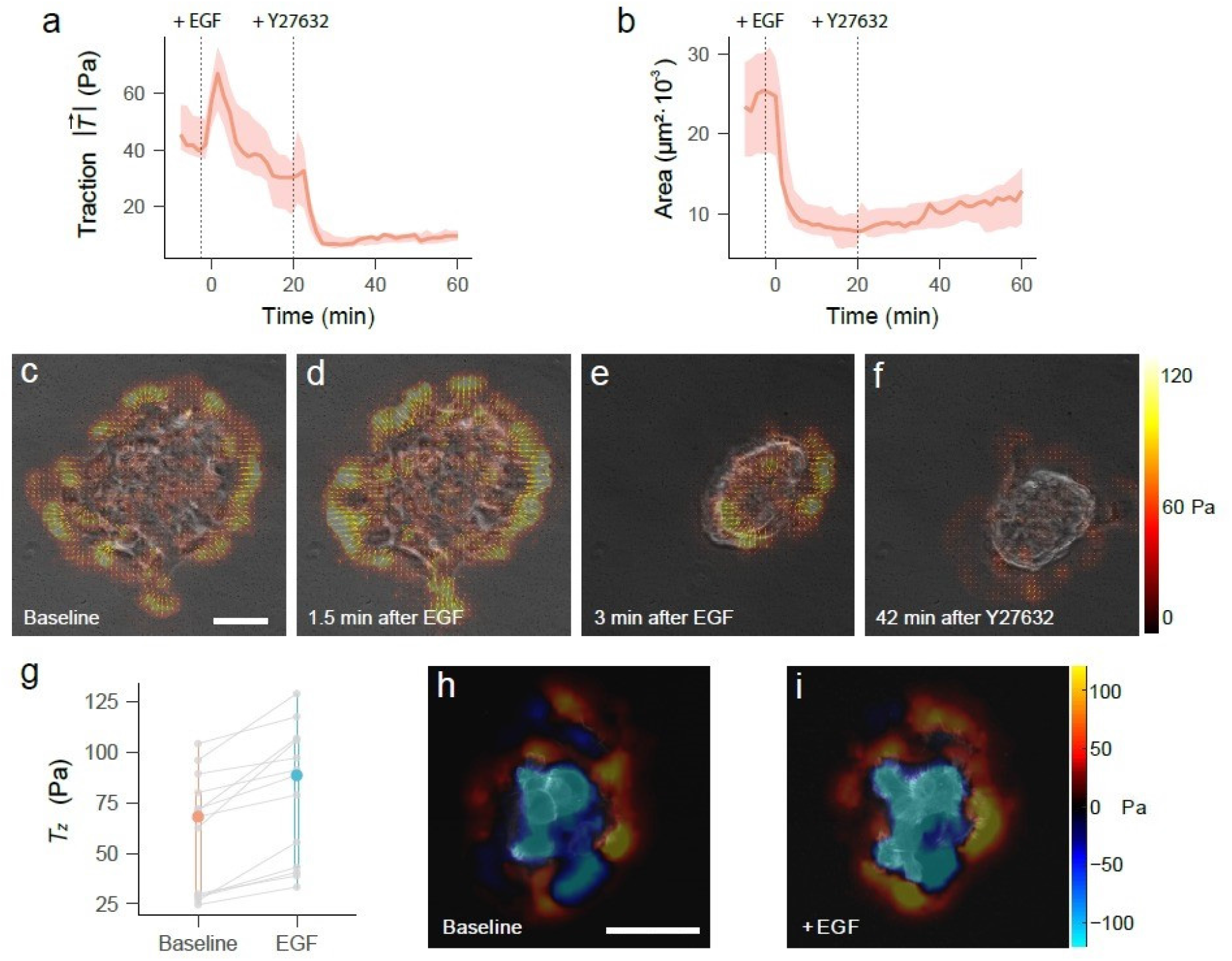
hEGF increases cell contractility and surface tension. **a,** Time lapse evolution of the modulus of the traction forces upon addition of 10 ng/ml of hEGF and the subsequent addition of 5μg/ml of Y27632 (indicated by vertical dashed lines). Gel stiffness is 6 kPa. **b,** Cluster area variation during the protocol described in (**a**). Data in panels **a-b** are median ± 95% CI estimated by bootstrapping (n=15 clusters). **c-f,** In-plane traction forces exerted by a representative cluster before addition of hEGF (**c**), 1.5 min after the addition of hEGF (**d**), 3 min after the addition of hEGF (**e**), and 42 min after the addition of Y27632 (**f**). **g,** The normal traction component *T_z_* increases 2 min after adding hEGF, indicating an increase in surface tension and Laplace pressure. A paired permutation test (two-tailed) indicated a significant increase (n= 13, p-value= 0.0002441). **h-i,** *T_z_* maps exerted by a representative cluster before (**h**) and after (**i**) the addition of hEGF.

**Extended Data Figure 7.**
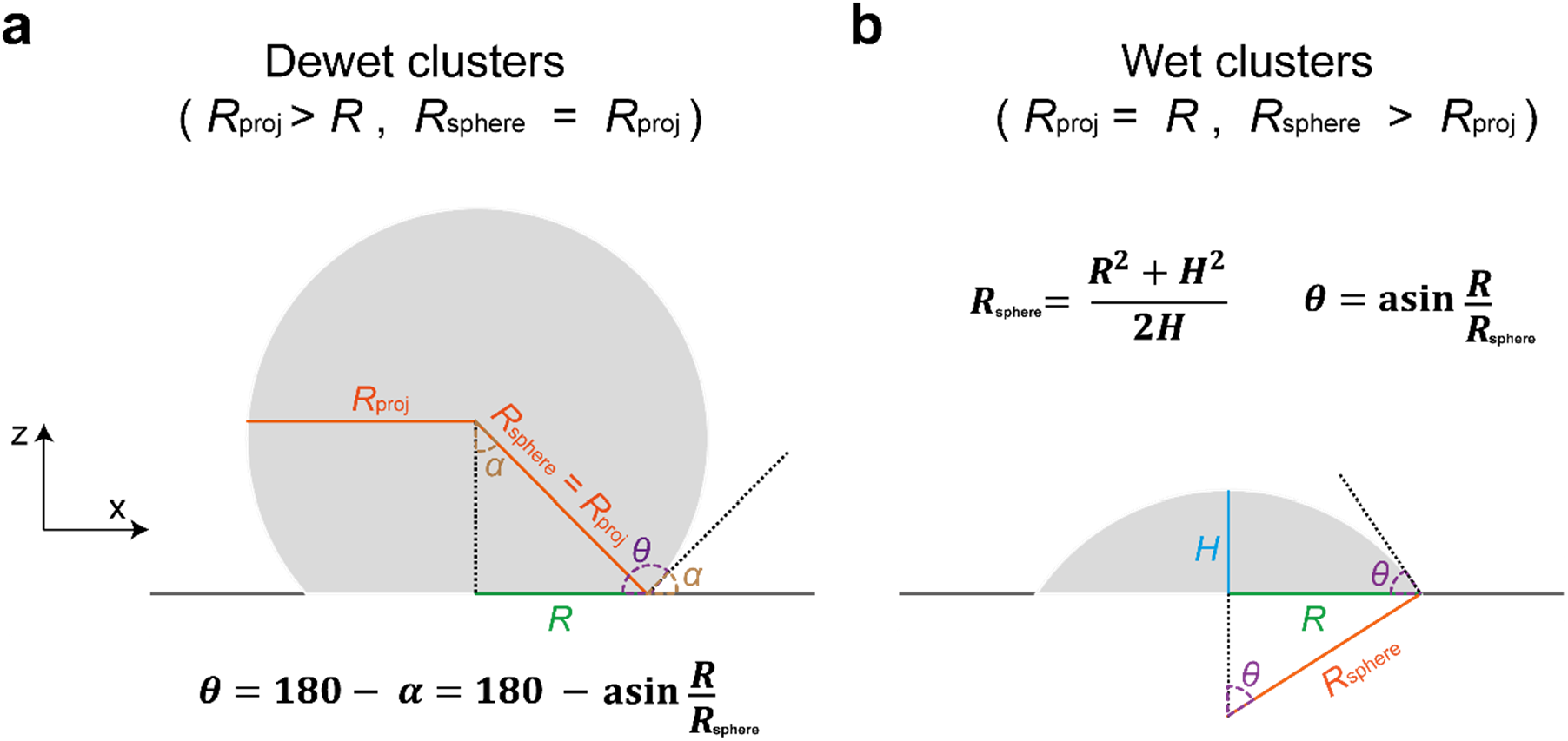
Calculation of contact angle. Measurement of contact angle *θ* as a function of the cluster radius (*R*_sphere_), the contact radius (*R*) and the cluster height (*H*). For low wettability (**a**) and high wettability (**b**) clusters the calculation varied. All parameters were estimated from high resolution z-stacks of LifeAct-mCherry A431 clusters seeded on E-cadherin substrates.

### Video legends

**Supplementary video 1**: Representative phase contrast video of A431 cell cluster migration on uniform stiffness gels of 0.2, 6, 24 and 200kPa coated with E-cadherin.

**Supplementary video 2:** z-stack movies of representative mCherry-Lifeact A431 cell clusters seeded on uniform stiffness gels of 0.2, 6, 24 and 200kPa coated with E-cadherin to illustrate cluster wetting state.

**Supplementary video 3:** Filopodia dynamics of representative A431 cell clusters seeded on uniform stiffness gels of 6, 24 and 200kPa coated with E-cadherin.

**Supplementary video 4:** Actin retrograde flow of representative A431 cell clusters seeded on uniform stiffness gels of 6, 24 and 200kPa coated with E-cadherin.

**Supplementary video 5:** Representative phase images of A431 cell cluster migration on a stiffness gradient coated with E-cadherin. Bottom numbers indicate stiffness in kPa.

**Supplementary video 6:** Migration of representative mCherry-Lifeact A341 cell clusters on a stiffness gradient (local stiffness = 10kPa) coated with E-cadherin.

**Supplementary video 7:** Representative phase contrast images of A431 cell cluster migration on a stiffness gradient coated with E-cadherin in controls and in presence of 0.5μM Y-27632 (ROCK inhibitor) aiming to partially inhibit cell contractility.

**Supplementary video 8:** Representative phase contrast images of A431 cell cluster migration on uniform stiffness gels of 0.2, 6, 24 and 200kPa coated with fibronectin in controls and in presence of 10ng/mL hEGF aiming to promote cell contractility.

**Supplementary video 9:** Representative phase contrast images of A431 cell cluster migration on a stiffness gradient coated with fibronectin in controls and in presence of 10ng/mL hEGF aiming to promote cell contractility.

**Supplementary video 10:** Phase contrast video representing an example of a sudden retraction of filopodia triggering a long durotactic hop.

## References

1. Haeger, A., Wolf, K., Zegers, M. M. & Friedl, P. Collective cell migration: guidance principles and hierarchies. Trends Cell Biol. 25, 556–566 (2015).

2. Majumdar, R., Sixt, M. & Parent, C. A. New paradigms in the establishment and maintenance of gradients during directed cell migration. Curr. Opin. Cell Biol. 30, 33–40 (2014).

3. Lyon, J. G., Carroll, S. L., Mokarram, N. & Bellamkonda, R. V. Electrotaxis of Glioblastoma and Medulloblastoma Spheroidal Aggregates. Sci. Rep. 9, 5309 (2019).

4. Lo, C. M., Wang, H. B., Dembo, M. & Wang, Y. L. Cell movement is guided by the rigidity of the substrate. Biophys. J. 79, 144–152 (2000).

5. Vincent, L. G., Choi, Y. S., Alonso-Latorre, B., del Álamo, J. C. & Engler, A. J. Mesenchymal stem cell durotaxis depends on substrate stiffness gradient strength. Biotechnol. J. 8, 472–484 (2013).

6. Sunyer, R. & Trepat, X. Durotaxis. Curr. Biol. 30, R383–R387 (2020).

7. Shellard, A. & Mayor, R. Durotaxis: The Hard Path from In Vitro to In Vivo. Dev. Cell 56, 227–239 (2021).

8. Zhu, M. et al. Spatial mapping of tissue properties in vivo reveals a 3D stiffness gradient in the mouse limb bud. Proc. Natl. Acad. Sci. 117, 4781–4791 (2020).

9. Shellard, A. & Mayor, R. Collective durotaxis along a self-generated stiffness gradient in vivo. Nature 600, 690–694 (2021).

10. Evans, N. D., Oreffo, R. O. C., Healy, E., Thurner, P. J. & Man, Y. H. Epithelial mechanobiology, skin wound healing, and the stem cell niche. J. Mech. Behav. Biomed. Mater. 28, 397–409 (2013).

11. DuChez, B. J., Doyle, A. D., Dimitriadis, E. K. & Yamada, K. M. Durotaxis by Human Cancer Cells. Biophys. J. 116, 670–683 (2019).

12. Sunyer, R. et al. Collective cell durotaxis emerges from long-range intercellular force transmission. Science 353, 1157 (2016).

13. Martinez, J. S., Schlenoff, J. B. & Keller, T. C. S. Collective epithelial cell sheet adhesion and migration on polyelectrolyte multilayers with uniform and gradients of compliance. Exp. Cell Res. 346, 17–29 (2016).

14. Alert, R. & Casademunt, J. Role of Substrate Stiffness in Tissue Spreading: Wetting Transition and Tissue Durotaxis. Langmuir 35, 7571–7577 (2019).

15. Pi-Jaumà, I., Alert, R. & Casademunt, J. Collective durotaxis of cohesive cell clusters on a stiffness gradient. Eur. Phys. J. E 45, 7 (2022).

16. Escribano, J. et al. A hybrid computational model for collective cell durotaxis. in Biomech. model. mechanobiol. 17, 1037–1052 (2018).

17. Novikova, E. A., Raab, M., Discher, D. E. & Storm, C. Persistence-driven durotaxis: Generic, directed motility in rigidity gradients. Phys. Rev. Lett. 118, 078103 (2017).

18. Isomursu, A. et al. Directed cell migration towards softer environments. Nat. Mater. 1–10 (2022).

19. Lazopoulos, K. A. & Stamenović, D. Durotaxis as an elastic stability phenomenon. J. Biomech. 41, 1289–1294 (2008).

20. Yu, G., Feng, J., Man, H. & Levine, H. Phenomenological modeling of durotaxis. Phys. Rev. E 96, 010402 (2017).

21. Rens, E. G. & Merks, R. M. H. Cell Shape and Durotaxis Explained from Cell-Extracellular Matrix Forces and Focal Adhesion Dynamics. iScience 23, 101488 (2020).

22. Isenberg, B. C., DiMilla, P. A., Walker, M., Kim, S. & Wong, J. Y. Vascular Smooth Muscle Cell Durotaxis Depends on Substrate Stiffness Gradient Strength. Biophys J 97, 1313–1322 (2009).

23. Hartman, C. D., Isenberg, B. C., Chua, S. G. & Wong, J. Y. Vascular smooth muscle cell durotaxis depends on extracellular matrix composition. Proc. Natl. Acad. Sci. 113, 11190–11195 (2016).

24. Richardson, B. E. & Lehmann, R. Mechanisms guiding primordial germ cell migration: strategies from different organisms. Nat. Rev. Mol. Cell Biol. 11, 37–49 (2010).

25. Cai, D. et al. Mechanical feedback through E-cadherin promotes direction sensing during collective cell migration. Cell 157, 1146–1159 (2014).

26. Dai, W. et al. Tissue topography steers migrating Drosophila border cells. Science 370, 987–990 (2020).

27. Grimaldi, C. et al. E-cadherin focuses protrusion formation at the front of migrating cells by impeding actin flow. Nat. Commun. 11, 5397 (2020).

28. Dorrell, M., Aguilar, E. & Friedlander, M. Dorrell, M. I., Aguilar, E. & Friedlander, M. Retinal vascular development is mediated by endothelial filopodia, a preexisting astrocytic template and specific R-cadherin adhesion. Invest. Ophthalmol. Vis. Sci. 43, 3500-3510. Invest. Ophthalmol. Vis. Sci. 43, 3500–10 (2002).

29. Luccardini, C. et al. N-cadherin sustains motility and polarity of future cortical interneurons during tangential migration. J. Neurosci. Off. J. Soc. Neurosci. 33, 18149–18160 (2013).

30. Padmanaban, V. et al. E-cadherin is required for metastasis in multiple models of breast cancer. Nature 573, 439–444 (2019).

31. Pérez-González, C. et al. Active wetting of epithelial tissues. Nat. Phys. 15, 79–88 (2019).

32. Douezan, S. et al. Spreading dynamics and wetting transition of cellular aggregates. Proc. Natl. Acad. Sci. U. S. A. 108, 7315–7320 (2011).

33. Douezan, S., Dumond, J. & Brochard-Wyart, F. Wetting transitions of cellular aggregates induced by substrate rigidity. Soft Matter 8, 4578–4583 (2012).

34. Gonzalez-Rodriguez David, Guevorkian Karine, Douezan Stéphane, & Brochard-Wyart Françoise. Soft Matter Models of Developing Tissues and Tumors. Science 338, 910–917 (2012).

35. Beaune, G. et al. How cells flow in the spreading of cellular aggregates. Proc. Natl. Acad. Sci. 111, 8055–8060 (2014).

36. Wallmeyer, B., Trinschek, S., Yigit, S., Thiele, U. & Betz, T. Collective Cell Migration in Embryogenesis Follows the Laws of Wetting. Biophys. J. 114, 213–222 (2018).

37. Beaune, G. et al. Spontaneous migration of cellular aggregates from giant keratocytes to running spheroids. Proc. Natl. Acad. Sci. U. S. A. 115, 12926–12931 (2018).

38. Alert, R. & Trepat, X. Physical Models of Collective Cell Migration. Annu. Rev. Condens. Matter Phys. 11, 77–101 (2020).

39. Riveline, D. et al. Focal contacts as mechanosensors: externally applied local mechanical force induces growth of focal contacts by an mDia1-dependent and ROCK-independent mechanism. J. Cell Biol. 153, 1175–1186 (2001).

40. Ghibaudo, M. et al. Traction forces and rigidity sensing regulate cell functions. Soft Matter 4, 1836–1843 (2008).

41. Elosegui-Artola, A. et al. Mechanical regulation of a molecular clutch defines force transmission and transduction in response to matrix rigidity. Nat. Cell Biol. 18, 540–548 (2016).

42. Barry, A. K. et al. α-Catenin cytomechanics – role in cadherin-dependent adhesion and mechanotransduction. J. Cell Sci. 127, 1779–1791 (2014).

43. Sunyer, R., Jin, A. J., Nossal, R. & Sackett, D. L. Fabrication of Hydrogels with Steep Stiffness Gradients for Studying Cell Mechanical Response. PLoS ONE 7, e46107 (2012).

44. Blanch-Mercader, C. et al. Effective viscosity and dynamics of spreading epithelia: a solvable model. Soft Matter 13, 1235–1243 (2017).

45. Walcott, S. & Sun, S. X. A mechanical model of actin stress fiber formation and substrate elasticity sensing in adherent cells. Proc. Natl. Acad. Sci. 107, 7757–7762 (2010).

46. Saez, A. et al. Traction forces exerted by epithelial cell sheets. J. Phys. Condens. Matter 22, 194119 (2010).

47. Marcq, P., Yoshinaga, N. & Prost, J. Rigidity sensing explained by active matter theory. Biophys. J. 101, L33–35 (2011).

48. Trichet, L. et al. Evidence of a large-scale mechanosensing mechanism for cellular adaptation to substrate stiffness. Proc. Natl. Acad. Sci. 109, 6933–6938 (2012).

49. Sens, P. Rigidity sensing by stochastic sliding friction. Europhys. Lett. 104, 38003 (2013).

50. Gupta, M. et al. Adaptive rheology and ordering of cell cytoskeleton govern matrix rigidity sensing. Nat. Commun. 6, 7525 (2015).

51. Guevorkian, K., Colbert, M.-J., Durth, M., Dufour, S. & Brochard-Wyart, F. Aspiration of biological viscoelastic drops. Phys. Rev. Lett. 104, 218101 (2010).

52. Guevorkian, K., Gonzalez-Rodriguez, D., Carlier, C., Dufour, S. & Brochard-Wyart, F. Mechanosensitive shivering of model tissues under controlled aspiration. Proc. Natl. Acad. Sci. 108, 13387–13392 (2011).

53. Manning, M. L., Foty, R. A., Steinberg, M. S. & Schoetz, E.-M. Coaction of intercellular adhesion and cortical tension specifies tissue surface tension. Proc. Natl. Acad. Sci. 107, 12517–12522 (2010).

54. Chan, G. K., McGrath, J. A. & Parsons, M. Spatial activation of ezrin by epidermal growth factor receptor and focal adhesion kinase co-ordinates epithelial cell migration. Open Biol. 11, 210166 (2021).

55. Iwabu, A., Smith, K., Allen, F. D., Lauffenburger, D. A. & Wells, A. Epidermal growth factor induces fibroblast contractility and motility via a protein kinase C delta-dependent pathway. J. Biol. Chem. 279, 14551–14560 (2004).

56. Czirók, A., Schlett, K., Madarász, E. & Vicsek, T. Exponential Distribution of Locomotion Activity in Cell Cultures. Phys. Rev. Lett. 81, 3038–3041 (1998).

57. Wu, P.-H., Giri, A., Sun, S. X. & Wirtz, D. Three-dimensional cell migration does not follow a random walk. Proc. Natl. Acad. Sci. 111, 3949–3954 (2014).

58. González-Valverde, I. & García-Aznar, J. M. Mechanical modeling of collective cell migration: An agent-based and continuum material approach. Comput. Methods Appl. Mech. Eng. 337, 246–262 (2018).

59. Garcia-Gonzalez, D. & Muñoz-Barrutia, A. Computational insights into the influence of substrate stiffness on collective cell migration. Extreme Mech. Lett. 40, 100928 (2020).

60. Deng, Y., Levine, H., Mao, X. & Sander, L. M. Collective Motility, Mechanical Waves, and Durotaxis in Cell Clusters. ArXiv:2007.10488 [physics.bio-ph] (2020).

61. Yousafzai, M. S. et al. Active Regulation of Pressure and Volume Defines an Energetic Constraint on the Size of Cell Aggregates. Phys. Rev. Lett. 128, 048103 (2022).

62. Yousafzai, M. S. et al. Tissue pressure and cell traction compensate to drive robust aggregate spreading. bioRxiv:2020.08.29.273334 (2020).

63. Style, R. W. et al. Patterning droplets with durotaxis. Proc. Natl. Acad. Sci. 110, 12541–12544 (2013).

64. Style, R. W., Jagota, A., Hui, C.-Y. & Dufresne, E. R. Elastocapillarity: Surface Tension and the Mechanics of Soft Solids. Annu. Rev. Condens. Matter Phys. 8, 99–118 (2017).

65. Babb, S. G. & Marrs, J. A. E-cadherin regulates cell movements and tissue formation in early zebrafish embryos. Dev. Dyn. 230, 263–277 (2004).

66. Shimizu, T. et al. E-cadherin is required for gastrulation cell movements in zebrafish. Mech. Dev. 122, 747–763 (2005).

67. Shamir, E. R. et al. Twist1-induced dissemination preserves epithelial identity and requires E-cadherin. J. Cell Biol. 204, 839–856 (2014).

68. Giannone, G., Mège, R.-M. & Thoumine, O. Multi-level molecular clutches in motile cell processes. Trends Cell Biol. 19, 475–486 (2009).

69. Nguyen, T. et al. Enhanced cell–cell contact stability and decreased N-cadherin-mediated migration upon fibroblast growth factor receptor-N-cadherin cross talk. Oncogene 38, 6283–6300 (2019).

70. Rakshit, S., Zhang, Y., Manibog, K., Shafraz, O. & Sivasankar, S. Ideal, catch, and slip bonds in cadherin adhesion. Proc. Natl. Acad. Sci. 109, 18815–18820 (2012).

71. Changede, R. & Sheetz, M. Integrin and cadherin clusters: A robust way to organize adhesions for cell mechanics. BioEssays 39, e201600123 (2017).

72. Bays, J. L. et al. Vinculin phosphorylation differentially regulates mechanotransduction at cell–cell and cell–matrix adhesions. J. Cell Biol. 205, 251–263 (2014).

73. Sehgal, P. et al. Epidermal growth factor receptor and integrins control force-dependent vinculin recruitment to E-Cadherin junctions. J Cell Sci 15, 131 (2018).

74. Tewari, S. et al. Statistics of shear-induced rearrangements in a two-dimensional model foam. Phys. Rev. E 60, 4385–4396 (1999).

75. Ben-Zion, Y. & Rice, J. R. Slip patterns and earthquake populations along different classes of faults in elastic solids. J. Geophys. Res. Solid Earth 100, 12959–12983 (1995).

76. Sethna, J. P., Dahmen, K. A. & Myers, C. R. Crackling noise. Nature 410, 242–250 (2001).

77. Barriga, E. H., Franze, K., Charras, G. & Mayor, R. Tissue stiffening coordinates morphogenesis by triggering collective cell migration in vivo. Nature 554, 523–527 (2018).

78. Tiscornia, G., Singer, O. & Verma, I. M. Production and purification of lentiviral vectors. Nat. Protoc. 1, 241–245 (2006).

79. Chevalier, S. et al. Creating Biomimetic Surfaces through Covalent and Oriented Binding of Proteins. Langmuir 26, 14707–14715 (2010).

80. Gräslund, S. et al. Protein production and purification. Nat. Methods 5, 135–146 (2008).

81. Trepat, X. et al. Physical forces during collective cell migration. Nat. Phys. 5, 426–430 (2009).

82. Rico, F. et al. Probing mechanical properties of living cells by atomic force microscopy with blunted pyramidal cantilever tips. Phys. Rev. E 72, 021914 (2005).

83. Alcaraz, J. et al. Microrheology of Human Lung Epithelial Cells Measured by Atomic Force Microscopy. Biophys. J. 84, 2071–2079 (2003).

