## Supplementary Note for "Stiffness-dependent active wetting enables optimal collective cell durotaxis"

Here, we present a theoretical framework to explain collective durotaxis of cell clusters in terms of a continuum model of the tissue. To this end, we describe a cell cluster as an active fluid droplet that partially wets the substrate. Extending a previous 2D model of tissue wetting, we first develop a theory of 3D active wetting for tissues on a uniform substrate. In this theory, the contact line dynamics results from the balance between the in-plane forces acting at the basal monolayer and the out-of-plane surface tension of the cell cluster, defining a dynamic contact angle. We then apply this theory to predict cluster durotaxis on a substrate stiffness gradient, explaining the generic non-monotonic dependence of the durotactic velocity with substrate stiffness observed in our experiments. Our analysis shows that the existence of an optimal stiffness for collective durotaxis is connected to the wetting properties of the droplet, which depend on stiffness. In particular we show that the maximum of durotactic velocity occurs near the point where the contact angle crosses over 90 degrees. We then study how the durotactic velocity varies with the mechanical parameters of the tissue, which allows us to explain the dependence of durotaxis on tissue size and cellular contractility observed in our experiments.

#### 1 Theory of 3D active wetting

We describe a cell cluster on a substrate as an active liquid droplet that partially wets a solid surface with a contact angle $\theta$ (Fig. 3a). The interface between the cell cluster and the surrounding passive fluid has a surface tension $\gamma$, which results from a combination of passive cell-cell adhesion and active cortical tension [1–5]. Averaging out tissue shape fluctuations, we describe the cell cluster as a spherical cap of radius $R$. We assume that the bulk of the cell cluster is passive, and that the dynamics of the droplet is determined by the interplay of its surface tension and the in-plane forces at the basal cell monolayer. Following previous work [6], we model the basal cell monolayer as a 2D active polar fluid. We then extend this model to 3D by proposing a generalized Young-Dupré force balance that includes the out-of-plane contribution of surface tension. This new ingredient modifies the contact line dynamics and defines the droplet's dynamic contact angle. Then, taking active cellular forces that depend on substrate stiffness [7–12], we predict how this environmental property affects the droplet's contact angle and spreading dynamics. Finally, we use our theory to show how these stiffness-dependent wetting properties give rise to collective durotaxis of cell clusters on stiffness gradients.

##### 1.1 2D model of the basal cell monolayer

Following Refs. [6, 13–15], we model the basal cell monolayer as a 2D active polar fluid described by two fields, cell polarity $\boldsymbol{p}(\boldsymbol{r},t)$ and velocity $\boldsymbol{v}(\boldsymbol{r},t)$. We assume that the polarity field is independent of the flow field, since the timescale of cell repolarization through contact inhibition of locomotion ($\tau_{\mathrm{CIL}} \sim 10$ min [16, 17]) is much smaller than the typical shear rate of flows in the monolayer ($\tau \sim 100$ min [18, 19]). Thus, the polarity field follows a purely relaxational dynamics, $\partial_t p_\alpha \propto -\delta F/\delta p_\alpha$. The tendency of cells to align with their neighbors is encoded in the effective free energy

$$F = \int \left[ \frac{a}{2} p_\alpha p_\alpha + \frac{K}{2} (\partial_\alpha p_\beta)(\partial_\alpha p_\beta) \right] \mathrm{d}^2 \boldsymbol{r}, \tag{S1}$$

being $K$ the Frank constant that defines the energetic cost of the polarity gradients [20], and $a > 0$ a restoring coefficient that favors the unpolarized state $p = 0$ in the bulk. We further assume a quasistatic evolution of the polarity $\partial_t p_\alpha = 0$, hence

$$L_c^2 \nabla^2 p_\alpha = p_\alpha, \tag{S2}$$

where $L_c \equiv \sqrt{K/a}$ is the nematic length that defines the width of a boundary layer in which cells are polarized at the periphery of the monolayer. Since cells at the edge are polarized toward free space, we impose the boundary condition $p_\alpha = n_\alpha$, where $n_\alpha$ is the outward normal vector. Solving Eq. (S2) with this boundary condition, the modulus of the polarity decays from one at the edge to zero in the bulk over the characteristic length $L_c$.

We describe tissue flow based on the force balance equation

$$\partial_\beta \sigma_{\alpha\beta} + f_\alpha = 0, \tag{S3}$$

where $\sigma_{\alpha\beta}$ is the stress tensor, and $f_\alpha$ is the external force density originated at the tissue-substrate interface. Note that the experimentally measured monolayer tension and traction are $\sigma_{\alpha\beta} h$ and $T_\alpha \equiv -f_\alpha h$, respectively, being $h$ the height of the monolayer. In Eq. (S3) we have neglected inertia, consistently with the small values of Reynolds number for tissue flows. We have also neglected pressure gradients by assuming that the 2D fluid is highly compressible as in-plane compression and expansion are accommodated by local changes in the monolayer thickness, without significant changes in pressure [21].

As tissue spreading takes place over very long time scales, we neglect the short-time elastic response of the tissue and assume a purely viscous response [21]. Hence, we take the constitutive equation for the stress tensor of a compressible active viscous polar fluid [22–26]. This includes two main contributions: active contractile stresses originated in cell-cell forces and quantified by the contractility parameter $\zeta < 0$, and viscous stresses that result from cell-cell adhesion forces and are weighted by an effective viscosity $\eta$. Similarly, the cell-substrate force density $f_\alpha$ has two main contributions: an active traction force that is proportional to polarity $\boldsymbol{p}$ and with a maximum value $\zeta_i$, and a friction force due to cell-substrate adhesion, proportional to the tissue velocity $\boldsymbol{v}$ and with a coefficient $\xi$ [6, 27]. Altogether,

$$\sigma_{\alpha\beta} = \eta(\partial_\alpha v_\beta + \partial_\beta v_\alpha) - \zeta p_\alpha p_\beta, \tag{S4}$$

$$f_\alpha = -\xi v_\alpha + \zeta_i p_\alpha. \tag{S5}$$

To complete the model we need to specify the boundary conditions for the stress. This is the point where the effect of the 3D shape of cell cluster will be incorporated, so we defer this discussion to Section 1.3.

#### 1.2 The 2D active wetting transition for cell monolayers

The 2D model of tissue spreading presented above predicts an active wetting transition in cell monolayers, which was verified in experiments [6]. This transition has no analogue in the classical theory of wetting; it results from the competition between active traction that favors tissue spreading and contractile stresses that favor retraction. The transition is controlled by a characteristic length of active polar fluids, defined as the ratio of contractile nematic stresses to active polar forces: $L_p \equiv |\zeta|/\zeta_i$. The active wetting transition takes place at a critical value $L_p^*$: the tissue spreads for $L_p < L_p^*$ (wetting), and it retracts for $L_p > L_p^*$ (dewetting).

The physical properties of this active wetting transition depend on whether dissipation is dominated by the internal tissue viscosity $\eta$ or by the external friction $\xi$. The comparison of these two effects defines the so-called screening length $\lambda \equiv \sqrt{\eta/\xi}$, which indicates an effective range of hydrodynamic interactions. The limit $\lambda \to \infty$ is known as the 'wet' limit, for which hydrodynamic interactions are long-ranged; the limit $\lambda \to 0$ is known as the 'dry' limit, for which hydrodynamic interactions are screened out by friction.

The study of 2D active wetting in Ref. [6] considered circular monolayers in the 'wet' limit. In that case, $L_p^* \sim R$, with $R$ being the monolayer radius. Therefore, the wetting transition occurs at a critical radius $R^*$: larger monolayers $R > R^*$ spread, and smaller monolayers $R < R^*$ retract. The existence of a critical radius illustrates the non-local character of active wetting in the 'wet' limit. In contrast to classic wetting theory, the advance or recession of the tissue front is not determined by local forces at the interface, but by the system as a whole. More recent works have considered the 'dry' limit, when friction forces are dominant over viscosity [13–15]. In this case, $L_p^* \sim \lambda$ [15], so there is no critical size for the wetting-dewetting transition. In this limit, the advance or recession of the front is determined by the local forces at the interface, and hence the wetting transition is size-independent.

Here, we will address the generic case and keep both friction and viscosity. For simplicity, we will solve the active wetting problem considering only one dimension ($\hat{\boldsymbol{x}}$), along which the cell monolayer spreads or retracts (Fig. 3a). This corresponds to assuming translational invariance along the perpendicular direction. The predictions for the active wetting transition in this one-dimensional geometry differ only in geometrical factors from those for circular cell monolayers [6, 13–15]. In addition, the one-dimensional geometry is particularly suited when tissue migration is biased by external gradients, such as in the experiments on collective durotaxis with rectangular cell monolayers [28]. It also yields a good approximation for the migration of circular monolayers as tissue flows occur mainly in the direction of the imposed gradient [15]. In the one-dimensional geometry, Eqs. (S3) to (S5) combine into

$$2\eta\partial_x^2 v = 2\zeta p \partial_x p + \xi v - \zeta_i p, \tag{S6}$$

where

$$p(x) = \frac{\sinh\left((x - X)/L_c\right)}{\sinh\left(R/L_c\right)} \tag{S7}$$

is the solution of Eq. (S2), with $R$ being the monolayer half-width (radius), and $X$ being the center-of-mass position. Given $x_+$ and $x_-$ the positions of the two edges of the monolayer, we have $X \equiv (x_+ + x_-)/2$ and $R \equiv (x_+ - x_-)/2$. With the solution of the velocity profile $v(x)$ at a given time, the velocity of the center-of-mass $v_X$ and the spreading velocity $v_S$ are given by

$$v_X \equiv \dot{X} = \frac{v(x_+) + v(x_-)}{2}, \quad v_S \equiv \dot{R} = \frac{v(x_+) - v(x_-)}{2}. \tag{S8}$$

In the absence of an external gradient, we have $v_X = 0$. However, $v_S$ may be positive when the monolayer is spreading (wetting), or negative when it is retracting (dewetting). The 2D wetting transition is thus defined by $v_S = 0$.

For reference, we quote the results for the spreading velocity $v_S$ in both the wet and the dry limits. Assuming $L_c \ll R, \lambda$, and with stress-free boundary conditions at the monolayer edge, we have [15]

$$v_S^{\text{wet}} \approx \frac{L_c \zeta_i}{2\eta}\left(R - \frac{L_p}{2}\right), \tag{S9}$$

$$v_S^{\text{dry}} \approx \frac{L_c \zeta_i}{2\eta}\left(\sqrt{2}\lambda - \frac{L_p}{2}\right). \tag{S10}$$

#### 1.3 Generalized Young-Dupré equilibrium for 3D cell clusters

Here, we extend the theory of active wetting to 3D cell clusters. To this end, we add the contribution of the out-of-plane surface tension $\gamma$ of the cell cluster to the 2D model of the basal cell monolayer presented above (Fig. 3a). As discussed, for stress-free boundary conditions, a cell monolayer either spreads or retracts indefinitely as a result of the competition between active traction and contractility. However, for 3D droplets, the surface tension $\gamma$ introduces an additional force that can either favor spreading or retraction depending on the contact angle $\theta$. For $\theta > 90°$, surface tension favors spreading (Fig. 3a left), whereas for $\theta < 90°$, surface tension favors retraction (Fig. 3a right). For sufficiently large surface tension, and depending on parameter values, all three active forces may balance and the droplet may reach a partial wetting state, namely a stable equilibrium with a finite contact angle $\theta$ (Figs. S1 and S2).

In classical wetting theory, the equilibrium contact angle is determined by the Young-Dupré condition, which establishes the balance of the three surface tensions at the contact line. For cell aggregates, a similar

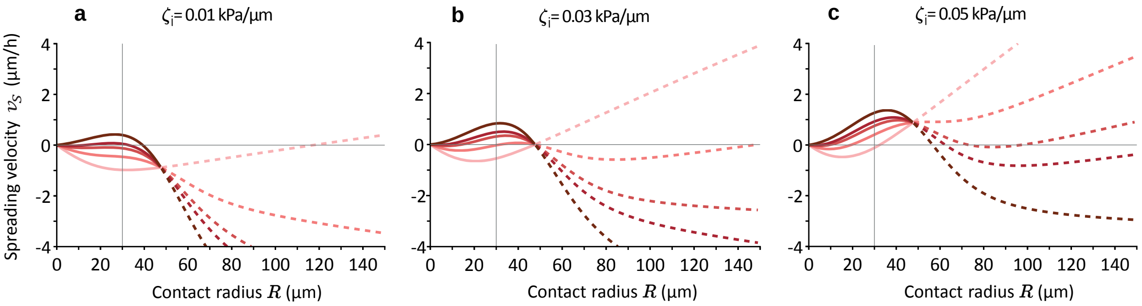


**Figure S1. Roots of the spreading velocity define the partial wetting state for sufficiently large surface tension** $\gamma$. Spreading velocity of a cluster on top of a uniform-stiffness substrate as a function of the contact radius $R$, for different values of active traction (**a-c**) and surface tension $\gamma = 0, 1.5, 2.5, 3, 4$ mN/m (from lighter to darker red). The other parameter values are those from Table SI, with a constant friction $\xi = 0.22$ kPa s/µm$^2$. The volume is kept constant to $V = 2/3\pi R^{*3}$, being $R^* = 47.6$ µm the radius at the 2D wetting transition for $\zeta_i = 0.03$ kPa/µm (**b**). We switch from solid to dashed lines when the cluster has a contact angle of 90° ($R = R_{\text{sphere}} = H$), and so the continuous lines correspond to angles $\theta > 90°$ ($R < R^*$) and the dashed lines to angles $\theta < 90°$ ($R > R^*$). The vertical line at each panel is drawn at $R = 30$ µm, which is the initial size ($R_0$) chosen for the dynamics in Fig. S2. From this point, we can predict whether a cluster will converge to the stable fixed point ($v_S(R_0) > 0$ with roots), or will completely dewet ($v_S(R_0) < 0$) or wet the substrate ($v_S(R_0) > 0$ without roots).

energetic approach was proposed to define the wetting conditions in terms of the cell-cell and cell-substrate adhesion energies [29–32]. This approach, however, does not explicitly account for active cellular forces, which have a key role in tissue wetting [6]. Following Ref. [6], here we assume that the surface tension of the interface between the surrounding passive fluid and the substrate is negligible in front of the active forces. Consistently, we propose a generalized Young-Dupré condition that captures a balance of three active forces: active traction and contractility, which are distributed in the polarized layer of the basal cell monolayer, and the cluster's surface tension, which enters as a local force at the contact line. Hence, we include the horizontal component of surface tension into the monolayer force balance Eq. (S6) as a stress boundary condition:

$$n_\alpha (h\sigma_{\alpha\beta}) n_\beta = -\gamma \cos\theta, \tag{S11}$$

where $n_\beta$ is the unit normal vector at the monolayer edge, and $h\sigma_{\alpha\beta}$ is the monolayer tension, with $h$ the monolayer height. In turn, the vertical component of the surface tension is balanced by the Young-Laplace pressure $P = 2\gamma/R_{\text{sphere}}$, which the cell cluster exerts on the substrate. This relationship allows us to infer $\gamma$ from measurements of vertical traction forces (Fig. 1g-j), which provide a direct measurement of $P$ (see table 2 in Methods).

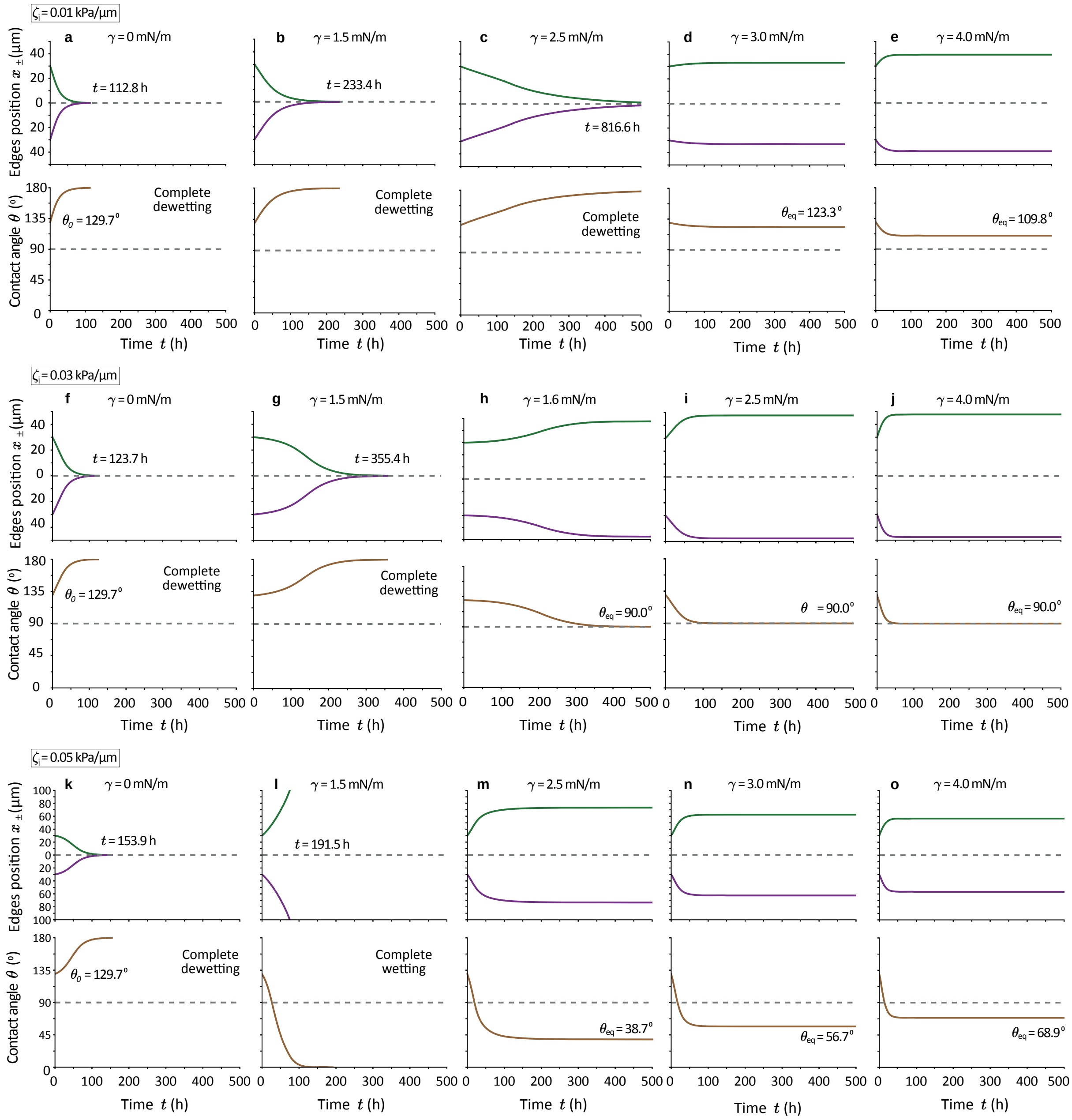

**Figure S2. Cell clusters reach an equilibrium state of partial wetting for sufficiently large surface tension $\gamma$.** Evolution of the position of the edges of the basal cell monolayer (green for right $x_+$, and purple for left $x_-$) and the contact angle of a cluster on top of a uniform-stiffness substrate. We show the evolution for three different values of the active traction (**a-e**, **f-j** and **k-o**), and for different values of surface tension, increasing from left to right. In all cases, the initial contact radius is $R_0 = 30$ µm and the height $H_0 = 63.9$ µm, giving a volume (which is kept constant throughout the evolution) of $V = 2/3\pi R^{*3}$, being $R^* = 47.6$ µm the radius at the 2D wetting transition for $\zeta_i = 0.03$ kPa/µm. The parameters are those from Table SI, with $\xi = 0.22$ kPa s/µm$^2$, a simulation time $T = 500$h and a time step $\Delta t = 6$ min. **a-e**, For $\gamma = 0$ mN/m (**a**), the cluster fully dewets the substrate (defined when it reaches $R \leq 0.1$ µm) after 112.8h. Increasing surface tension to $\gamma = 1.5, 2.5, 3, 4$ mN/m (**b**,**c**,**d**,**e**), the cluster reaches the equilibrium contact angle only if $\gamma$ is large enough, with $\theta_{\text{eq}} = 123.3, 109.8^\circ$ (**d** and **e**). **f-j**, For $\gamma = 0$ mN/m (**f**), the cluster fully dewets the substrate after 123.7h. With $\gamma = 1.5, 1.6, 2.5, 4$ mN/m (**g**,**h**,**i**,**j**), the cluster reaches the equilibrium contact angle of $\theta_{\text{eq}} = 90^\circ$ only if $\gamma$ is large enough. **k-o**, For $\gamma = 0$ mN/m (**k**), the cluster fully wets the substrate (defined when it reaches $H \leq 0.1$ µm) after 153.9h. With $\gamma = 1.5, 2.5, 3, 4$ mN/m (**l**,**m**,**n**,**o**), the cluster reaches the equilibrium contact angle only if $\gamma$ is large enough, with $\theta_{\text{eq}} = 38.7, 56.7, 68.9^\circ$. The larger $\gamma$, the closer is $\theta_{\text{eq}}$ to $90^\circ$.

| | Description | Typical value |
|---|---|---|
| $h$ | Monolayer height | 5 µm [6, 33] |
| $L_c$ | Nematic length | 15 µm [6, 18] |
| $\eta$ | Monolayer viscosity | 20 MPa s [6, 18] |
| $\zeta$ | Monolayer contractility | $-2$ kPa [6] |
| $\lambda$ | Hydrodynamic screening length | $4.2 \times 10^2$ µm |
| $E_0$ | Substrate's softest stiffness | 0.5 kPa (Fig. 2a) |
| $E'$ | Stiffness gradient | 33 kPa/mm (Fig. 2a) |
| $E^*$ | Characteristic stiffness of force saturation | 140 kPa [31] |
| $\zeta_i^\infty$ | Traction saturation value | 0.3 kPa/µm [6, 18] |
| $\xi^\infty$ | Friction saturation value | 0.5 kPa s/µm$^2$ [34] |
| $\xi_0$ | Friction at $E \to 0$ | $2.2 \times 10^{-1}$ kPa s/µm$^2$ |
| $P_{0x}$ | Pressure at $E_0$ | 4.2 Pa |
| $s_P$ | Pressure sensitivity to stiffness | $1.8 \times 10^{-2}$ |

**Table SI.** Symbols and typical values of model parameters, of the same order of magnitude as in the references.

### 2 Collective durotaxis of 3D cell clusters

#### 2.1 Stiffness-dependent forces

In this section, we study collective durotaxis of cell clusters by imposing a gradient of substrate stiffness. Following previous work [14, 15], we assume that the parameters that encode tissue-substrate interactions depend on the local substrate stiffness. This functional dependence must be determined independently and is an input into our hydrodynamic model. From Refs. [7–12], we assume that both friction and traction parameters saturate to maximal values at high stiffness, and hence we take

$$\zeta_i(E) = \zeta_i^\infty \frac{E}{E + E^*}, \quad \xi(E) = \xi^\infty \frac{E}{E + E^*} + \xi_0, \tag{S12}$$

where $\zeta_i^\infty$ and $\xi^\infty$ are saturation values, and $E^*$ is a characteristic stiffness of force saturation. Finally, $\xi_0$ is the friction coefficient at vanishing substrate stiffness, which we add to avoid the strict 'wet' limit $\lambda \to \infty$ ($\xi \to 0$). This limit is ill-defined in presence of a traction gradient, because global force balance cannot be satisfied in the absence of friction[1].

In our experimental measurements, not only radial in-plane tractions (Fig. 1g-i) but also out-of-plane tractions (Fig. 1g,h,j) increase with substrate stiffness. We model this mechanosensitive response of the tissue surface tension by making the pressure $P(E)$ increase with substrate stiffness. For simplicity, and because there is no experimental evidence, to our knowledge, of saturation of out-of-plane tractions with stiffness, we assume a linear dependence

$$P(E) = P_0 + s_P E. \tag{S13}$$

We call $s_P$ the pressure sensitivity to stiffness. Via the Young-Laplace relation, the pressure dependence on stiffness corresponds to a stiffness-dependent surface tension

$$\gamma(E) = \frac{R_{\text{sphere}}}{2} P(E) = \gamma_0 + \ell_\gamma E, \tag{S14}$$

where $\gamma_0 = P_0 R_{\text{sphere}}/2$ is the bare surface tension, and we call $\ell_\gamma = s_P R_{\text{sphere}}/2$ the stiffness response length. Note that when the radius of the spherical cap $R_{\text{sphere}}$ varies, as in dynamical evolutions with constant droplet volume $V$, $\gamma_0$ and $\ell_\gamma$ also vary accordingly.

[1] To see this, consider the total traction, i.e., the cell-substrate force density given by Eq. (S5): $\boldsymbol{f} = -\xi \boldsymbol{v} + \zeta_i \boldsymbol{p}$. In the absence of friction, we have $\boldsymbol{f} = \zeta_i \boldsymbol{p}$, whose integral over the tissue, $\int \boldsymbol{f}\, \mathrm{d}^2\boldsymbol{r}$, does not vanish if the traction is different on the stiff and the soft edges.

#### 2.2 Durotaxis of 2D cell monolayers

Before analyzing the full 3D problem, we discuss the durotactic migration of a 2D cell monolayer. To this end, we take a linear stiffness profile

$$E(x) = E_0 + E'x, \tag{S15}$$

being $E_0$ and $E'$ the stiffness offset and gradient, respectively. Introducing the stiffness profile in Eqs. (S13) and (S14), we obtain the pressure and surface tension profiles

$$P(x) = P_{0x} + P'x = (P_0 + s_P E_0) + s_P E'x \tag{S16}$$

$$\gamma(x) = \gamma_{0x} + \gamma'x = (\gamma_0 + \ell_\gamma E_0) + \ell_\gamma E'x. \tag{S17}$$

Furthermore, to reduce the number of parameters and gain physical insight, we momentarily replace the saturating traction and friction coefficients given in Eq. (S12) by non-saturating linear functions of substrate stiffness $E$, or equivalently, of the position $x$ along the stiffness gradient:

$$\zeta_i(x) = \zeta_i^0 + \zeta_i'x, \quad \xi(x) = \xi_0 + \xi'x, \tag{S18}$$

where $\zeta_i'$ and $\xi'$ are the traction and friction gradients, respectively. We further simplify the problem by taking a uniform friction ($\xi' = 0$). In this case, Eq. (S6) can be solved analytically to obtain the velocity field, which was done Appendix B of Ref. [15]. In the dry and wet limits, and assuming $L_c \ll R, \lambda$, the center-of-mass velocity or durotactic velocity $v_X$ does not depend on the boundary conditions of the stress, and we obtain

$$v_X^{\text{dry}} \approx \frac{L_c\lambda}{2\eta} R\zeta_i' = \frac{L_c\lambda}{4\eta}(\zeta_i^+ - \zeta_i^-), \tag{S19}$$

$$v_X^{\text{wet}} \approx \frac{L_c\lambda^2}{2\eta}\zeta_i' = \frac{L_c\lambda^2}{4\eta}\frac{\zeta_i^+ - \zeta_i^-}{R}, \tag{S20}$$

where $\zeta_i^\pm$ are the local values of the active traction at the respective edges. In both cases, the spreading velocity $v_S$ has the same expression as in uniform-stiffness situations (Eqs. (S9) and (S10)) albeit replacing $\zeta_i$ by $\zeta_i(X)$, i.e. the active traction at the center-of-mass $X$.

In the dry limit, the spreading dynamics is local in the sense that the two edges move independently from the other, driven only by local forces at the edges. Thus, the durotactic velocity is directly proportional to the traction difference across the tissue, and hence to the monolayer size (Eq. (S19)). In contrast, in the wet limit the two edges are hydrodynamically coupled, and the durotactic velocity depends directly on the traction difference and inversely on the monolayer size $R$ (Eq. (S20)). These two dependencies cancel each other, and the durotactic velocity is independent of the monolayer size in this case (Eq. (S20)).

In this simplest situation with a uniform traction gradient $\zeta_i'$ and uniform friction ($\xi' = 0$), the durotactic velocity of a cell monolayer is independent of the traction offset $\zeta_i^0$, and hence of the local substrate stiffness [15]. This behavior cannot explain our experimental measurements, which show a non-monotonic dependence of durotactic velocity with substrate stiffness (Fig. 2e,f).

#### 2.3 Non-monotonic durotaxis of 3D cell clusters

In this section we explain the non-monotonic dependence of durotactic velocity with substrate stiffness measured in our experiments (Fig. 2e,f). We show that this non-monotonic behavior arises from a combination of different effects: a) Both the increase of friction and the saturation of traction with stiffness produce a decrease of durotactic velocity at high stiffness, and b) the variation of the contact angle, which is controlled by the out-of-plane surface tension, produces an increase of durotactic velocity at low stiffness. The competition of these opposite trends yields a maximal durotactic velocity at an intermediate stiffness, for which durotaxis is optimal. We also show that this maximum occurs near the point where the contact angle crosses over 90°, which separates the conditions of low and high wettability. We call this region the neutral wetting regime.

To show this, we use the model to predict the evolution of the velocity and the shape of a cluster as it migrates toward stiffer regions, numerically integrating the dynamics of a cluster with constant volume (see Methods). A representative result is shown in Fig. 3b,c, which we discuss in the Main Text.

We remark that in our present model the wetting properties are defined globally for the cluster as a whole, and so we cannot contemplate contact angle differences at both edges due to different stiffness. However, the dominant asymmetry that drives durotaxis is the difference between tractions at both edges [15], so a contact angle asymmetry would enter only as a higher order correction. We will study this contribution in future work, including shape variations and deformability of the substrate.

##### 2.3.1 Slowdown of durotaxis at high stiffness

First, we focus on the decrease of durotactic velocity at high stiffness, which we explain based on the dynamics of the 2D basal cell monolayer. We start by analyzing the role of the friction coefficient $\xi$. As it increases with substrate stiffness, the durotactic velocity decreases with substrate stiffness. As shown in Fig. S3, the stronger the friction gradient $\xi'$ (Eq. (S18)), the stronger the decay of durotactic velocity at high stiffness.

Next, we analyze the role of force saturation at high stiffnesses. As stiffness increases, active traction forces tend to saturate (Fig. S4a), and hence the traction difference across the tissue, $\Delta\zeta_i \equiv \zeta_i^+ - \zeta_i^-$, decreases (Fig. S5a, for a fixed contact radius $R$). Because the durotactic velocity increases with the traction difference $\Delta\zeta_i$ (Fig. S5b), traction saturation leads to a decrease of durotactic velocity with stiffness (Fig. S5c). As force saturation is pushed toward higher stiffness by increasing the crossover value $E^*$, the slowdown of durotaxis is less pronounced (Fig. S4b).

Altogether, both the increase in friction and the saturation of active traction lead to a slower durotaxis at high stiffnesses. Each of these two effects independently slows down durotaxis. Furthermore, both are independent of the 3D shape of the cluster; they arise even in the absence of tissue surface tension as shown in Fig. S5.

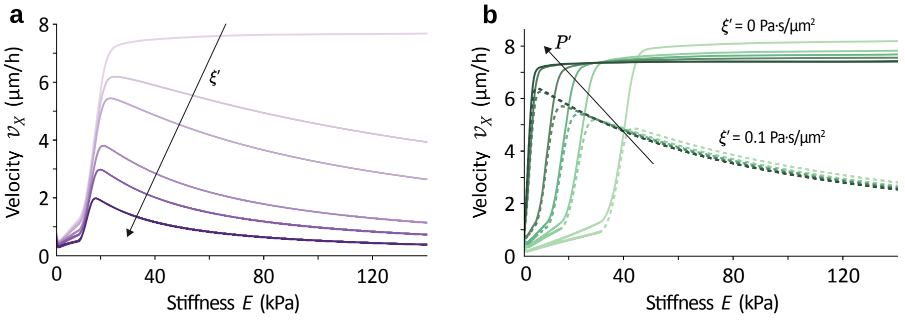


**Figure S3. Increasing friction produces a decrease of durotactic velocity with substrate stiffness.** Evolution of cluster motion as it moves along a stiffness gradient. The initial shape of the cluster is such with a contact radius $R_0 = 3.0$ μm and height $H_0 = 56.8$ μm, giving $\theta_0 = 174.0°$ (consistent to the experimental measurements of Fig. 1e). Thus, the volume is kept constant to $V = 92500\pi/3$ μm$^3$. The initial position is $x_0 = 100$ μm ($E_0 = 3.8$ kPa), and we choose linear traction, friction, and pressure profiles (Eqs. (S16) to (S18)) to show that force saturation is not required for the slowdown of durotaxis at high stiffness. **a**, An increasing friction gradient $\xi' = (0, 0.05, 0.1, 0.3, 0.5, 1.0)$ Pa s/μm$^2$ (from lighter to darker purple), lowers the durotactic velocity at high stiffness. **b**, An increasing pressure gradient $P' = (0.2, 0.4, 0.6, 1.0, 3.0, 4.0)$ Pa/μm (from lighter to darker green), yields smaller contact angles and hence faster durotaxis at low stiffness. However, pressure effects do not produce a velocity decrease at high stiffness; the decrease is only observed when a friction gradient is present (dashed lines). Other parameter values are listed in Table SI except for $\zeta_i^0 = 0.68$ Pa/μm and $\zeta_i' = 0.05$ Pa/μm$^2$, and the simulation time-step is $\Delta t = 6$ min.

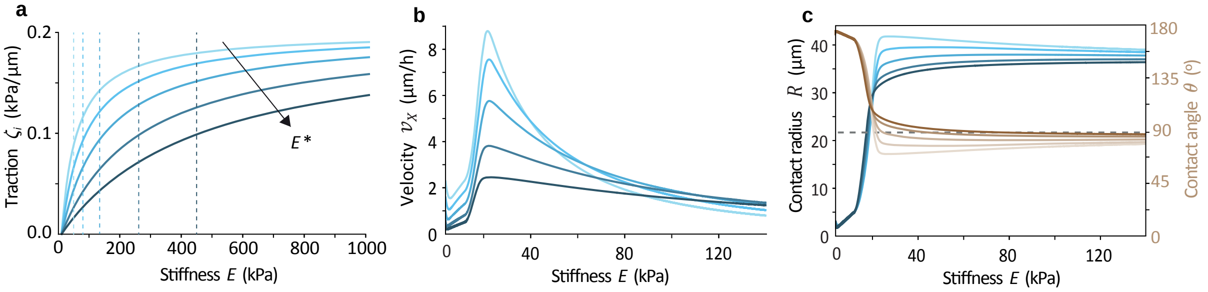


**Figure S4. Traction saturation produces a decrease of durotactic velocity with substrate stiffness. a,** Active traction profiles against stiffness values, whose saturation shifts to higher stiffness when the crossover stiffness $E^*$ increases (with $E^* = 50, 80, 140, 260, 450$ kPa, from lighter to darker blue). **b-c,** Evolution of cluster motion (**b**) and shape (**c**) as it moves along a stiffness gradient. As $E^*$ increases, the durotactic velocity and contact angle decrease are less pronounced. The initial shape of the cluster is such with a contact radius $R_0 = 3.0$ µm and height $H_0 = 56.8$ µm, giving $\theta_0 = 174.0°$ (consistent to the experimental measurements of Fig. 1e). Thus, the volume is kept constant to $V = 92500\pi/3$ µm$^3$, and the initial position is $x_0 = 100$ µm ($E_0 = 3.8$ kPa). Other parameter values are listed in Table SI, and the simulation time-step is $\Delta t = 6$ min. The slight increase of the contact angle for large stiffness values (and contact radius decrease), comes from the linear increase of the pressure with stiffness (Eq. (S13)), whereas traction and friction saturate. Thus, the surface tension (which is pointing inwards) is capable of causing monolayer retraction. The effect would be corrected if the pressure also saturated (as for instance in Fig. 3b,c) or decreased its value.

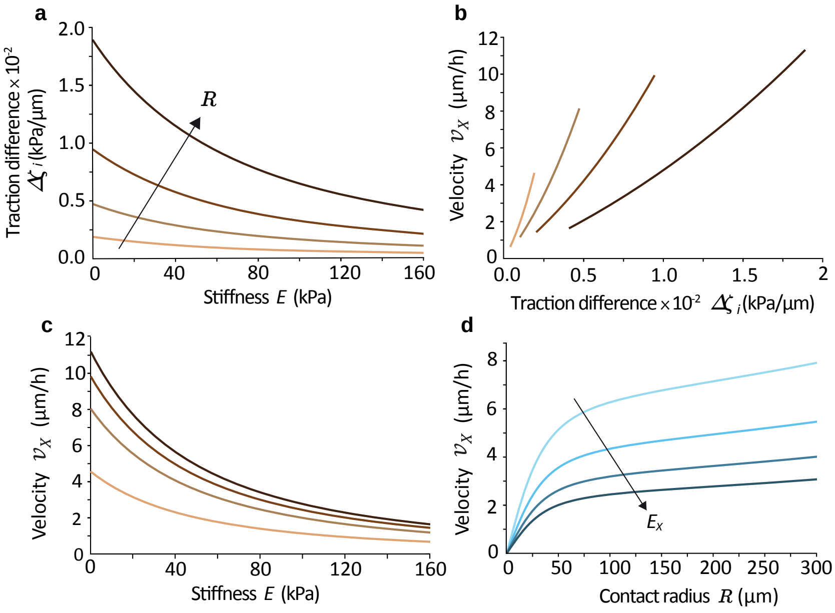


**Figure S5. Traction saturation produces a decrease of durotactic velocity with substrate stiffness and an increase with contact radius.** The figure corresponds to 2D monolayers, in the absence of tissue surface tension $\gamma = 0$. **a**, Because of traction saturation (Fig. S4a), the active traction difference across the monolayer $\Delta\zeta_i$ decreases with stiffness, but it increases with size $R$. In panels **a-c**, each curve is for a fixed contact radius $R$, with values $R = 20, 50, 100, 200$ µm from lighter to darker brown. **b-d**, The durotactic velocity increases with traction difference (**b**), and hence it decreases with stiffness (for a fixed $R$) (**c,d**). As the active traction difference is larger for larger $R$, the durotactic velocity increases with contact radius $R$ (for a fixed stiffness value $E$) (**c,d**). In panel **d**, each curve is for a fixed stiffness, with values $E_X = 25, 50, 75, 100$ kPa from lighter to darker blue. Other parameter values are listed in Table SI.

##### 2.3.2 Speed-up of durotaxis at low stiffness

Now, we focus on the increase of durotactic velocity at low stiffness, which can be explained based on how the 3D shape of the cluster changes with substrate stiffness. To this end, we use our 3D active wetting theory. From the experimental results (Fig. 1e,f), low stiffnesses promote cluster dewetting and hence lead to a high contact angle and low contact radius $R$. This small $R$ yields a small traction difference across the tissue, which implies a small durotactic velocity (Fig. S5c,d). Increasing the stiffness, the contact angle decreases and so $R$ increases (Fig. S4c and Fig. S6c), producing faster durotaxis. As it affects the contact angle, surface tension $\gamma$ -and thus pressure $P$- control the velocity increase, as shown in Fig. S3b and Fig. S6a,b.

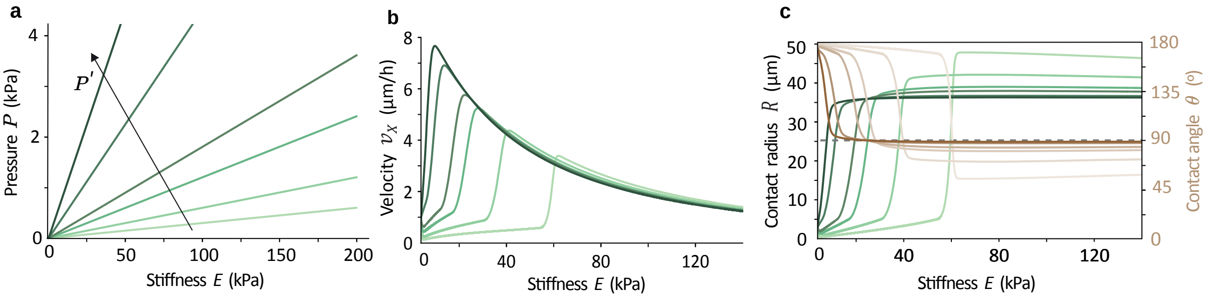


**Figure S6. Pressure increase controls the speed-up at low stiffness. a,** Pressure profiles against stiffness values, when the pressure gradient increases (with values $P' = (0.1, 0.2, 0.4, 0.6, 1.5, 3.0)$ Pa/µm, from lighter to darker green). **b-c,** Evolution of cluster motion (**b**) and shape (**c**) as it moves along a stiffness gradient. As $P'$ increases, the durotactic velocity and contact radius increase happens sooner, that is, for lower stiffness values. At high stiffnesses, the contact radius saturates and the durotactic velocity decreases due to the two other effects discussed in the text: friction increase (Fig. S3) and traction saturation (Fig. S4). The initial shape of the cluster is such with a contact radius $R_0 = 3.0$ µm and height $H_0 = 56.8$ µm, giving $\theta_0 = 174.0°$ (consistent to the experimental measurements of Fig. 1e). The volume is kept constant to $V = 92500\pi/3$ µm$^3$, and the initial position is $x_0 = 100$ µm ($E_0 = 3.8$ kPa). Other parameter values are listed in Table SI, and the simulation time-step is $\Delta t = 6$ min.

##### 2.3.3 Dynamic contact angle and location of optimal durotaxis

From the evolution of the clusters along the stiffness gradient, we find that the contact angle never coincides with an equilibrium solution at the corresponding stiffness (Fig. S1). The relaxation toward the equilibrium solution is slow compared to the time scale at which the cluster moves along the stiffness gradient. Accordingly, the observed contact angle is a dynamic one, and the evolution, except for extremely small stiffness gradients, is not quasistatic. Instead, as it performs durotaxis, the cluster has a non-zero spreading velocity, $v_S > 0$. This spreading increases the contact area of the cluster and therefore increases the durotactic velocity.

We see that $v_S(E)$ has a localized peak (blue curve in Fig. 3c and Figs. S9 to S11), which is related to the non-monotonic behavior of $v_S(R)$ in a uniform-stiffness substrate (Fig. S1). This peak gives place to the strongest speedup of the durotactic velocity. When $v_S$ decreases, the durotactic velocity increases more slowly, exhibiting an inflection point close to the location of the $v_S$ peak. Since this region of fast variations is correlated with fast variations of $\cos\theta$, which happen precisely at $\theta = 90°$, the maximal durotactic velocity will be typically near the crossing over 90° of the contact angle. Its exact location along the stiffness axis will depend on details of the profiles of traction and friction vs stiffness, which control the decrease of the durotactic velocity. For the type of profiles discussed here, the maximum of the durotactic velocity is indeed close to the peak of spreading velocity and the crossing over 90° of the contact angle. For extreme (and unrealistic) profiles (for instance for very small friction gradients), the location of maximal durotaxis may in principle depart significantly. Even in such cases though, the maximal values of durotactic velocity do not differ significantly from those reached past the peak of $v_S$, because the dependence of durotactic velocity with stiffness becomes very flat.

#### 2.4 The durotactic velocity depends on cluster size, cellular contractility, and stiffness gradient

Our experimental results (Fig. 2e,f,g) reveal that the durotactic velocity depends on the cluster size, cell contractility, and stiffness gradient. Here, we show how to capture all these results by modifying the corresponding parameters of our model.

##### 2.4.1 Cluster size

Our experiments show that larger clusters exhibit a higher durotactic velocity, which reaches its maximum at higher substrate stiffness (Fig. 2e). Our model recapitulates these two trends (Fig. 3e and Fig. S7a,d), which we explain in the following way: first, larger clusters have a larger traction difference across them, which drives faster durotaxis (Fig. S5c,d). Second, increasing cluster size favors monolayer wetting [6], and therefore larger clusters have lower contact angles. At low stiffness, the contact angle is always larger than 90°. In this regime, decreasing the contact angle, i.e. making it closer to 90°, implies that the horizontal component of the surface tension becomes smaller (Fig. 3a left). We have previously shown that the horizontal component of surface tension is responsible for the durotactic velocity increase at low stiffness (Fig. 3d, Fig. S6). Thus, larger clusters have a longer increase of durotactic velocity with substrate stiffness, and hence they reach their maximal velocity at higher stiffness.

##### 2.4.2 Cellular contractility

In our experiments, decreasing myosin-generated cellular contractility through the ROCK inhibitor Y-27632 produces slower durotaxis and shifts the maximum of the durotactic velocity toward lower stiffness (Fig. 2f). We recapitulate these trends with our model by decreasing the magnitude of all active forces, i.e. active traction $\zeta_i$, monolayer contractility $|\zeta|$, and tissue surface tension $\gamma$ (Fig. 3f and Fig. S7b,e). Specifically, we reduce active forces by multiplying their coefficients by factors $\alpha < 1$: $\zeta^{\text{red}} = \alpha_\zeta \zeta$, $\zeta_i^{\text{red}} = \alpha_{\zeta_i} \zeta_i$, $\gamma^{\text{red}} = \alpha_\gamma \gamma$.

Both monolayer contractility $\zeta$ and active traction $\zeta_i$ are active forces that would vanish completely with no myosin activity. In contrast, tissue surface tension $\gamma$ has both active and passive contributions [2]. The passive contribution is due to cell-cell adhesion, which would keep cells adhered and thus produce a tissue surface tension even without myosin activity. Therefore, a reduction of myosin activity, as induced by the Y-27632 treatment, affects $\zeta$ and $\zeta_i$ to a larger extent than $\gamma$. To account for this fact, we take $\alpha_\zeta = \alpha_{\zeta_i} < \alpha_\gamma$. Reducing active forces through these factors, we recover the decrease of durotactic velocity and the shift of its maximum toward lower stiffness.

Note that decreasing only the monolayer contractility $\zeta$ is not enough to explain the experimental results of the Y-27632 treatment. Decreasing only $|\zeta|$ promotes wetting [6], and therefore it yields higher contact radius (Fig. S8b), shifting the maximal durotactic velocity toward lower stiffness (Fig. S8a). This trend is consistent with the experimental results (Fig. 2f). However, decreasing only $|\zeta|$ also produces faster durotaxis (Fig. S8a), which is opposite to our experimental measurements. Thus, we confirm that to explain the results of the Y-27632 treatment, we need to implement a decrease of all three active forces as explained above.

##### 2.4.3 Stiffness gradient

Finally, our experiments show that a higher stiffness gradient produces faster durotaxis (Fig. 2g). Again, we recapitulate this result with our model (Fig. S7c,f). A higher stiffness gradient implies a larger traction difference across the basal monolayer, which drives faster durotaxis (Fig. S5b). In addition to explaining this result, our predictions reveal that the maximum of the durotactic velocity is shifted toward higher stiffness (Fig. S7c). This shift is due to the fact that a higher stiffness gradient yields a lower contact radius $R$ for a fixed $E$ (Fig. S7f), which we have previously shown to shift the optimal durotactic conditions to stiffer substrates (Fig. S6b,c).

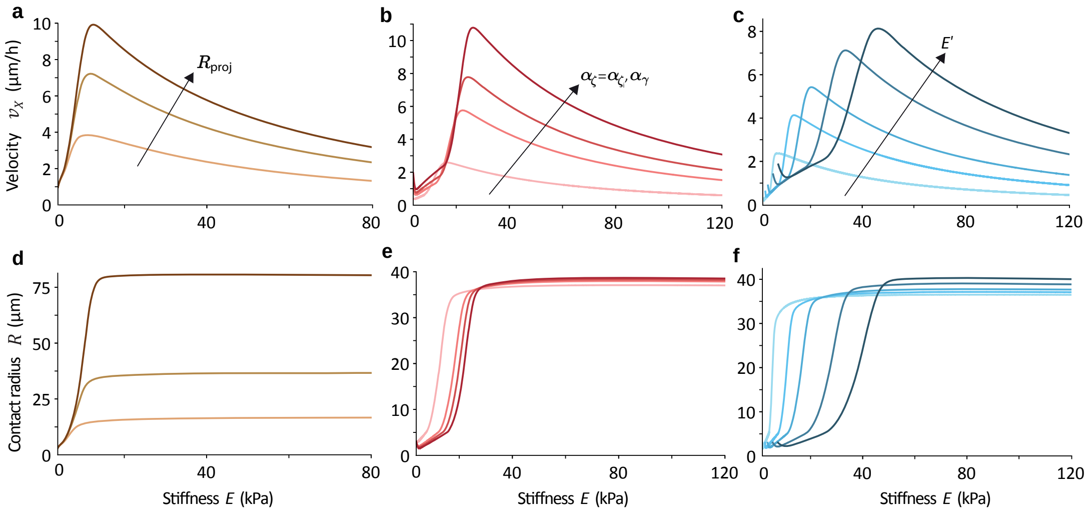


**Figure S7. The durotactic velocity depends on cluster size, cellular contractility, and stiffness gradient.** Evolution of cluster motion (**a-c**) and contact radius (**d-f**) as it moves along a stiffness gradient. **a**,**d**, Increasing the cluster's volume $V = 9250\pi/3, 92500\pi/3, 925000\pi/3$ µm$^3$, from lighter to darker brown, leads to larger contact radius (**d**), and hence faster durotaxis (**a**). **b**,**e**, Decreasing all the active forces through factors $\alpha_\zeta = \alpha_{\zeta_i} < \alpha_\gamma$ leads to slower durotaxis and a shift of the velocity maximum to smaller stiffnesses. We take $\alpha_\gamma = 0.7, 1, 1.2, 1.5$ and $\alpha_\zeta = \alpha_{\zeta_i} = 0.4, 1, 1.4, 2$ from lighter to darker red. A stronger decrease in cellular contractility corresponds to smaller $\alpha$. **c**,**f**, Increasing the stiffness gradient $E' = 10, 20, 30, 50, 70$ kPa/mm from lighter to darker blue, produces an increase of durotactic velocity and a displacement of its peak toward stiffer regions. In all the cases except from the lightest and darkest curves from **a** and **d**, the initial shape of the cluster is such with a contact radius $R_0 = 3.0$ µm and height $H_0 = 56.8$ µm, giving $\theta_0 = 174.0°$ (consistent to the experimental measurements of Fig. 1e). Thus, the volume is kept constant to $V = 92500\pi/3$ µm$^3$. In the two mentioned exceptions, $R_0 = 3.0$ µm and $H_0 = 26.1, 122.7$ µm, giving $\theta_0 = 166.9°, 177.2°$ respectively, and volume $V = 9250\pi/3, 925000\pi/3$ µm$^3$. The initial position is $x_0 = 100$ µm ($E_0 = 3.8$ kPa). Other parameter values are listed in Table SI, and the simulation time-step is $\Delta t = 6$ min.

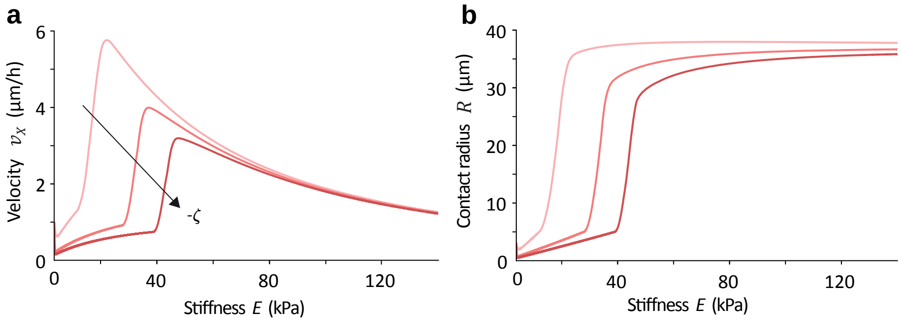


**Figure S8. Decreasing only monolayer contractility $|\zeta|$ does not capture the results of the Y-27632 treatment.** Evolution of cluster motion (**a**) and contact radius (**b**) as it moves along a stiffness gradient. Increasing monolayer contractility $-\zeta = 2, 5, 7$ kPa from lighter to darker red, produces slower durotaxis and shifts the maximal velocity to higher stiffness (**a**) as it promotes dewetting, i.e. smaller contact radius (**b**). The initial shape of the cluster is such with a contact radius $R_0 = 3.0$ µm and height $H_0 = 56.8$ µm, giving $\theta_0 = 174.0°$ (consistent to the experimental measurements of Fig. 1e). Thus, the volume is kept constant to $V = 92500\pi/3$ µm$^3$, and the initial position is $x_0 = 100$ µm ($E_0 = 3.8$ kPa). Other parameter values are listed in Table SI, and the simulation time-step is $\Delta t = 6$ min.

#### 2.5 Effects of other parameters in the dynamics of cell cluster durotaxis

Here, we analyze the effects of other parameters in the dynamical evolutions, such as the initial stiffness $E_0$ (Fig. S9), the contractility $|\zeta|$ (Fig. S10a,b), the pressure offset $P_0$ (Fig. S10c,d), and the pressure gradient $P'$ (Fig. S11).

We consider a cluster that starts with a high contact angle $\theta$ on a soft region of the substrate. As it advances toward stiffer regions, the cluster increases its wettability, lowering its contact angle $\theta$ and so expanding its contact radius $R$. As a result, the cluster increases its durotactic velocity; it speeds up. Eventually, when the cluster reaches sufficiently stiff substrates, the effects of friction increase and traction saturation become more important and the cluster slows down.

However, in many cases, there is an initial decrease of the durotactic velocity, corresponding to a decrease of the contact radius $R$. This initial slowdown happens when the cluster starts under conditions of dewetting, i.e. with a negative spreading velocity $v_S < 0$. In this situation, the contact radius initially decreases and the contact angle increases, as illustrated in Figs. S9 to S11, in which the initial contact angle is $\theta_0 = 136.4° > 90°$. In a softer initial position (Fig. S9), with higher contractility (Fig. S10a-c), or with smaller values of the pressure and thus the surface tension (Fig. S11), this early-stage effect is accentuated, since all these parameters favor cluster dewetting. Instead, a larger pressure diminishes this effect because it favors the expansion of $R$ at the initial stages and when $\theta > 90°$.

Finally, if the pressure does not increase with stiffness (Fig. S10d,e), surface tension offers a lower opposition to cluster spreading in low contact angle conditions ($\theta < 90°$). As a result, the contact radius keeps increasing significantly even at high stiffness). Nevertheless, the durotactic velocity still decreases due to the increase of friction forces.

Observing and characterizing this full spectrum of dynamical evolutions in experiments remains a challenge for future work.

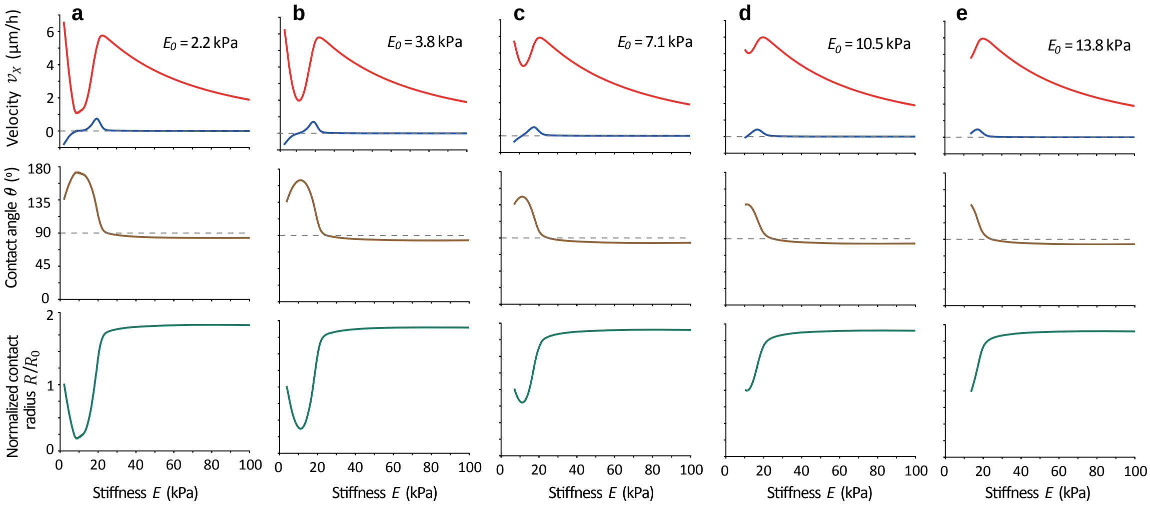


**Figure S9. Dynamics of cluster durotaxis when varying the initial substrate stiffness.** Evolution of cluster motion and shape as it moves along a stiffness gradient, with saturated traction and friction and linear pressure profiles. Regarding the velocity plots, the durotactic velocity is in red and the spreading velocity in blue. At each case, the initial shape of the cluster is such with a contact radius $R_0 = 20.0$ µm and height $H_0 = 50.0$ µm, giving $\theta_0 = 136.4°$ and a constant volume $V = 92500\pi/3$ µm$^3$. We change the initial position to $x_0 = 50, 100, 200, 300, 400$ µm, corresponding to $E_0 = 2.2, 3.8, 7.1, 10.5, 13.8$ kPa. Other parameter values are listed in Table SI, and the simulation time-step is $\Delta t = 6$ min.

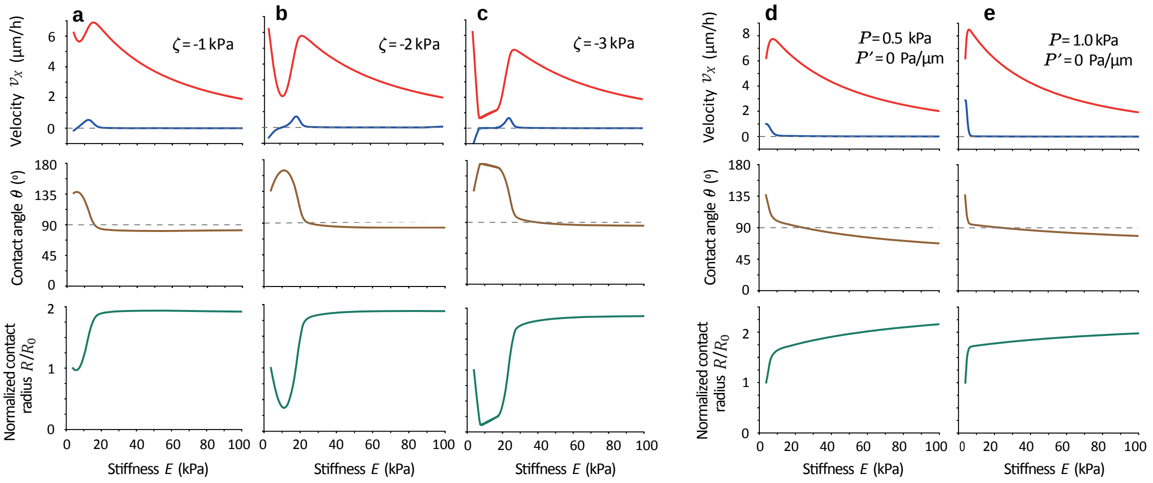


**Figure S10. Dynamics of cluster durotaxis when varying the contractility and pressure.** Evolution of cluster motion and shape as it moves along a stiffness gradient, with saturated traction and friction and linear pressure profiles. Regarding the velocity plots, the durotactic velocity is in red and the spreading velocity in blue. At each case, the initial shape of the cluster is such with a contact radius $R_0 = 20.0$ µm and height $H_0 = 50.0$ µm, giving $\theta_0 = 136.4^\circ$ and a constant volume $V = 92500\pi/3$ µm$^3$. The initial position is $x_0 = 100$ µm ($E_0 = 3.8$ kPa). In **a-c** we change the contractility to $-\zeta = 1, 2, 3$ kPa, with a linear pressure profile $P_{0x} = 4.2$ Pa and $P' = 0.6$ Pa/µm. In **d** and **e**, we take a uniform pressure of $P = 0.5$ and $1.0$ kPa respectively, with a fixed contractility $\zeta = -2$ kPa. Other parameter values are listed in Table SI, and the simulation time-step is $\Delta t = 6$ min.

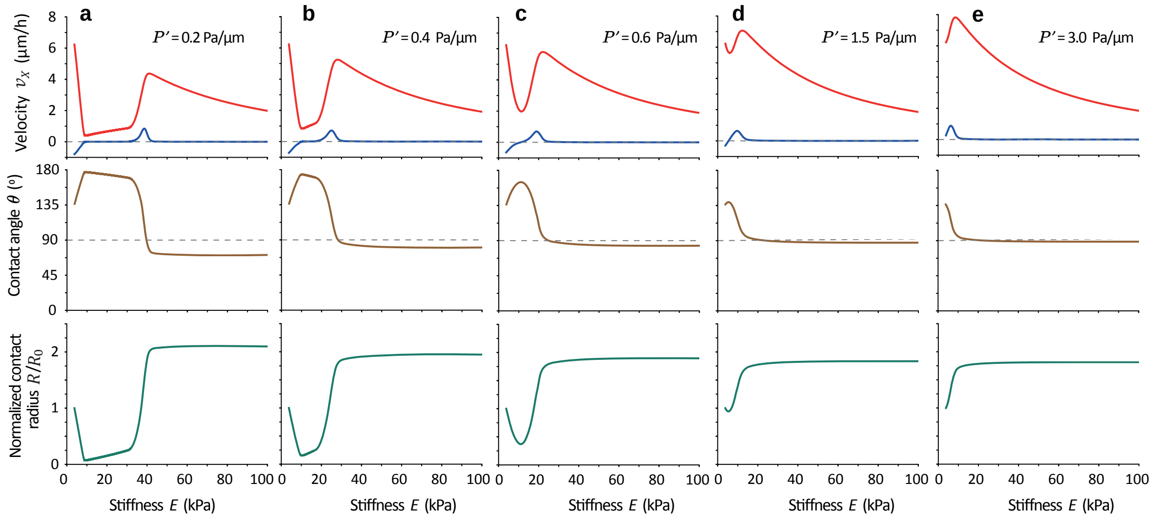


**Figure S11. Dynamics of cluster durotaxis when varying the pressure gradient.** Evolution of cluster motion and shape as it moves along a stiffness gradient, with saturated traction and friction and linear pressure profiles. Regarding the velocity plots, the durotactic velocity is in red and the spreading velocity in blue. At each case, the initial shape of the cluster is such with a contact radius $R_0 = 20.0$ µm and height $H_0 = 50.0$ µm, giving $\theta_0 = 136.4^\circ$ and a constant volume $V = 92500\pi/3$ µm$^3$. The initial position is $x_0 = 100$ µm ($E_0 = 3.8$ kPa), and we change the pressure gradient to $P' = (0.2, 0.4, 0.6, 1.5, 3.0)$ Pa/µm. Other parameter values are listed in Table SI, and the simulation time-step is $\Delta t = 6$ min.

### References


1. Lecuit, T. & Lenne, P. F. Cell surface mechanics and the control of cell shape, tissue patterns and morphogenesis. *Nat. Rev. Mol. Cell Biol.* **8,** 633–644 (2007).
2. Manning, M. L., Foty, R. A., Steinberg, M. S. & Schoetz, E.-M. Coaction of intercellular adhesion and cortical tension specifies tissue surface tension. *PNAS* **107,** 12517–12522 (2010).
3. Guevorkian, K., Colbert, M. J., Durth, M., Dufour, S. & Brochard-Wyart, F. Aspiration of biological viscoelastic drops. *Phys. Rev. Lett.* **104,** 218101 (2010).
4. Maître, J. L. & Heisenberg, C. P. The role of adhesion energy in controlling cell-cell contacts. *Curr. Opin. Cell Biol.* **23,** 508–514 (2011).
5. Ehrig, S. *et al.* Surface tension determines tissue shape and growth kinetics. *Sci. Adv.* **5,** 9394–9405 (2019).
6. Active wetting of epithelial tissues. *Nat. Phys.* **15,** 79–88 (2019).
7. Walcott, S. & Sun, S. X. A mechanical model of actin stress fiber formation and substrate elasticity sensing in adherent cells. *PNAS* **107,** 7757–7762 (2010).
8. Saez, A. *et al.* Traction forces exerted by epithelial cell sheets. *J. Phys.: Condens. Matter* **22,** 194119 (2010).
9. Trichet, L. *et al.* Evidence of a large-scale mechanosensing mechanism for cellular adaptation to substrate stiffness. *PNAS* **109,** 6933–6938 (2012).
10. Gupta, M. *et al.* Adaptive rheology and ordering of cell cytoskeleton govern matrix rigidity sensing. *Nat. Commun.* **6,** 7525 (2015).
11. Marcq, P., Yoshinaga, N. & Prost, J. Rigidity sensing explained by active matter theory. *Biophys. J.* **101,** L33–L35 (2011).
12. Sens, P. Rigidity sensing by stochastic sliding friction. *Europhys. Lett.* **104,** 38003 (2013).
13. Alert, R., Blanch-Mercader, C. & Casademunt, J. Active Fingering Instability in Tissue Spreading. *Phys. Rev. Lett.* **122,** 088104 (2019).
14. Alert, R. & Casademunt, J. Role of Substrate Stiffness in Tissue Spreading: Wetting Transition and Tissue Durotaxis. *Langmuir* **35,** 7571–7577 (2019).
15. Pi-Jaumà, I., Alert, R. & Casademunt, J. Collective durotaxis of cohesive cell clusters on a stiffness gradient. *Eur. Phys. J. E* **45,** 7 (2022).
16. Weber, G. F., Bjerke, M. A. & DeSimone, D. W. A Mechanoresponsive Cadherin-Keratin Complex Directs Polarized Protrusive Behavior and Collective Cell Migration. *Dev. Cell* **22,** 104–115 (2012).
17. Smeets, B. *et al.* Emergent structures and dynamics of cell colonies by contact inhibition of locomotion. *PNAS* **113,** 14621–14626 (2016).
18. Blanch-Mercader, C. *et al.* Effective viscosity and dynamics of spreading epithelia: a solvable model. *Soft Matter* **13,** 1235–1243 (2017).
19. Vincent, R. *et al.* Active Tensile Modulus of an Epithelial Monolayer. *Phys. Rev. Lett.* **115,** 248103 (2015).
20. De Gennes, P.-G. & Prost, J. *The Physics of Liquid Crystals* (Oxford, UK: Oxford Univ. Press. 2nd ed, 1993).
21. Alert, R. & Trepat, X. Physical Models of Collective Cell Migration. *Annu. Rev. Condens. Matter Phys.* **11,** 77–101 (2020).
22. Kruse, K., Joanny, J. F., Jülicher, F., Prost, J. & Sekimoto, K. Generic theory of active polar gels: A paradigm for cytoskeletal dynamics. *Eur. Phys. J. E* **16,** 5–16 (2005).
23. Jülicher, F., Kruse, K., Prost, J. & Joanny, J. F. Active behavior of the Cytoskeleton. *Phys. Rep.* **449,** 3–28 (2007).
24. Marchetti, M. *et al.* Hydrodynamics of soft active matter. *Rev. Mod. Phys.* **85,** 1143–1189 (2013).

25. Prost, J., Jülicher, F. & Joanny, J. F. Active gel physics. *Nat. Phys.* **11,** 111–117 (2015).
26. Jülicher, F., Grill, S. W. & Salbreux, G. Hydrodynamic theory of active matter. *Rep. Prog. Phys.* **81,** 076601 (2018).
27. Oriola, D., Alert, R. & Casademunt, J. Fluidization and Active Thinning by Molecular Kinetics in Active Gels. *Phys. Rev. Lett.* **118,** 088002 (2017).
28. Sunyer, R. *et al.* Collective cell durotaxis emerges from long-range intercellular force transmission. *Science* **353,** 1157–1161 (2016).
29. Ryan, P. L., Foty, R. A., Kohn, J. & Steinberg, M. S. Tissue spreading on implantable substrates is a competitive outcome of cell-cell vs. cell-substratum adhesivity. *PNAS* **98,** 4323–4327 (2001).
30. Douezan, S. *et al.* Spreading dynamics and wetting transition of cellular aggregates. *PNAS* **108,** 7315–7320 (2011).
31. Douezan, S., Dumond, J. & Brochard-Wyart, F. Wetting transitions of cellular aggregates induced by substrate rigidity. en. *Soft Matter* **8,** 4578–4583 (2012).
32. Beaune, G. *et al.* How cells flow in the spreading of cellular aggregates. *PNAS* **111,** 8055–8060 (2014).
33. Trepat, X. *et al.* Physical forces during collective cell migration. *Nat. Phys.* **5,** 426–430 (2009).
34. Cochet-Escartin, O., Ranft, J., Silberzan, P. & Marcq, P. Border forces and friction control epithelial closure dynamics. *Biophys. J.* **106,** 65–73 (2014).
